# Extracellular Ligand-Responsive Translational Regulation of Synthetic mRNAs using Engineered Receptors

**DOI:** 10.1101/2024.09.27.615322

**Authors:** Hideyuki Nakanishi, Keiji Itaka

## Abstract

mRNA drugs can encode any protein and are regarded as a promising therapeutic modality. In this study, we developed a new mRNA system allowing extracellular ligand-responsive translational regulation. This system consists of three mRNAs, two of which encode components for detecting extracellular ligands, and the other encodes the protein of interest, with a binding motif on 5’UTR for translational regulation. In the presence of ligand biomolecules such as arginine vasopressin (AVP) and prostaglandin E2 (PGE2), the translation level of the protein of interest is upregulated or downregulated in a ligand biomolecule concentration-dependent manner. We showed upregulation of anti-inflammatory signaling in response to an inflammatory mediator, PGE2, demonstrating that this system allowed production of the therapeutic proteins according to the disease site environment. This system will pave the way for the next-generation mRNA drugs that self-adjust their protein production levels in response to fluctuating disease states.

**Graphical Abstract:** 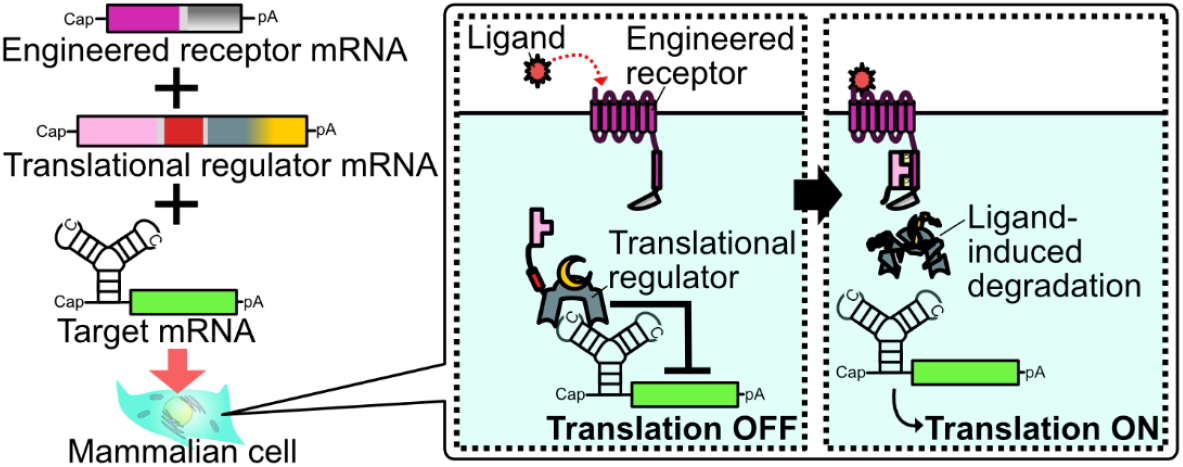

mRNAs encoding regulatory components (an engineered receptor and a translational regulator) and a target mRNA are co-transfected into mammalian cells. In the absence of the ligand, the target mRNA is translationally repressed by the translational regulator. The ligand induces engineered receptor-mediated degradation of the translational regulator protein, resulting in the translational activation of the target mRNA.

## Introduction

Synthetic mRNAs prepared via *in vitro* transcription (IVT) can express any protein without the risk of insertional mutagenesis^1^. Furthermore, synthetic mRNAs do not require nuclear import, thereby facilitating more efficient and uniform protein production than DNA-based vectors, such as plasmid DNA, especially in non-dividing cells^2^. Thus, synthetic mRNAs, such as mRNA drugs and mRNA vaccines, are regarded as promising therapeutic and prophylactic modalities. Despite their development in the early 1990s, synthetic mRNAs started to gain notable attention owing to the COVID-19 mRNA vaccines in 2020. They are currently being developed for a wide range of diseases such as cancers, monogenic disorders, and autoimmune diseases.^1^ Importantly, the therapeutic effects of mRNA drugs are not attributed to the mRNAs themselves but rather to the resulting translated proteins. Therefore, mRNA drugs have the potential to self-adjust their effects by regulating translation levels in response to disease states. Such self-adjustment can balance high therapeutic efficacy with minimal adverse effects by avoiding both insufficient and excessive therapeutic protein production.

Several groups, including ours, have developed translational regulation systems that respond to cell membrane-permeable molecules^3–8^, light irradiation^7,9,10^, and intracellular biomolecules such as RNAs^3,11–15^ and proteins^5–7,16–18^. While these systems enable cell-selective or manually inducible regulation, an extracellular ligand-responsive translational regulation system is more desirable for responding to disease states at the organ, tissue, or systemic levels, as disease-related cells secrete various disease markers such as hormones and lipid mediators. Thus, in the present study, we developed an extracellular ligand-responsive translational regulation system by combining an engineered receptor protein and a translational regulator protein. As this system can be designed to respond to various disease markers by selecting an appropriate receptor module, it is expected to provide disease state-matched therapeutic effects for various diseases.

## Materials and Methods

### Reagents and instruments

Reagents and instruments used in this study are listed below.

- D-MEM (High Glucose) with L-Glutamine, Phenol Red, and Sodium Pyruvate (FUJIFILM Wako Pure Chemical Corporation, Osaka, Japan, #043-30085)
- D-PBS(−) (FUJIFILM Wako Pure Chemical Corporation, Osaka, Japan, #049-29793)
- Fetal Bovine Serum (BioWest, Nuaillé, France, # S1400-500)
- Penicillin–streptomycin (Sigma-Aldrich Japan, Tokyo, Japan, #P4458)
- 0.25% trypsin/EDTA (Sigma-Aldrich Japan, Tokyo, Japan, #T4049)
- PrimeSTAR Max DNA polymerase (Takara Bio, Shiga, Japan, #R045A)
- KOD One PCR Master Mix -Blue-(Toyobo Co., Ltd., Osaka, Japan, #KMM-201)
- Custom DNA Oligos (Eurofins Genomics K.K., Tokyo, Japan)
- Monarch PCR & DNA Cleanup Kit (New England Biolabs Japan Inc., Tokyo, Japan, #T1030)
- In-Fusion HD Cloning Kit (Takara Bio, Shiga, Japan, # 639648)
- In-Fusion Snap Assembly Master Mix (Takara Bio, Shiga, Japan, # 638947)
- LB Broth (Lennox) (Sigma-Aldrich Japan, Tokyo, Japan, #L3022)
- LB Broth with agar (Lennox) (Sigma-Aldrich Japan, Tokyo, Japan, #L2897)
- Monarch Plasmid Miniprep Kit (New England Biolabs Japan Inc., Tokyo, Japan, #T1010)
- PureYield Plasmid Miniprep System (Promega K.K., Tokyo, Japan, #A1222).
- NanoDrop One (Thermo Fisher Scientific K.K., Kanagawa, Japan, #ND-ONE-W).
- MEGAscript T7 Transcription Kit (Thermo Fisher Scientific K.K., Kanagawa, Japan, #AMB13345)
- Takara IVTpro T7 mRNA Synthesis Kit (Takara Bio, Shiga, Japan, #6144)
- CleanCap Reagent AG (3’ OMe) (TriLink Biotechnologies, San Diego, CA, USA, #N-7413)
- N^1^-methylpseudouridine-5’-Triphosphate (TriLink Biotechnologies, San Diego, CA, USA, #N-1081)
- G(5’)ppp(5’)A RNA Cap Structure Analog (New England Biolabs Japan Inc., Tokyo, Japan, #S1406)
- RNAClean XP (Beckman Coulter Inc., Brea, CA, USA, #A63987)
- Quick CIP (New England Biolabs Japan Inc., Tokyo, Japan, #M0525)
- RNeasy Mini Kit (Qiagen K.K., Tokyo, Japan, #74104)
- Monarch RNA Cleanup Kit (New England Biolabs Japan Inc., Tokyo, Japan, #T2040)
- Qubit RNA Broad Range Assay Kit (Thermo Fisher Scientific K.K., Kanagawa, Japan, #Q10210)
- Qubit 4 Fluorometer (Thermo Fisher Scientific K.K., Kanagawa, Japan, #Q33238)
- Agilent 2100 Bioanalyzer (Agilent Technologies Japan Ltd., Tokyo, Japan)
- Agilent RNA6000 Nano Kit (Agilent Technologies Japan Ltd., Tokyo, Japan, #5067-1511)
- CELLSTAR μClear 96-well TC F-bottom (chimney well) white microplates (Greiner Bio-One International GmbH, Kremsmünster, Austria, #655098)
- Corning 96-well Flat Clear Bottom White Polystyrene TC-treated Microplates (Corning Inc., NY, USA, #3610)
- Lipofectamine MessengerMAX (Thermo Fisher Scientific K.K. Kanagawa, Japan, #LMRNA015)
- [Arg^8^]-vasopressin solution (Sigma-Aldrich Japan, Tokyo, Japan, #V0377)
- prostaglandin E2 (Tokyo Chemical Industry Co., Ltd., Tokyo, Japan, #P1884)
- bradykinin (FUJIFILM Wako Pure Chemical Corporation, Osaka, Japan, #029-19404)
- Nano-Glo Dual-Luciferase Assay System (Promega K.K., Tokyo, Japan, #N1630)
- GloMax Navigator Microplate Luminometer (Promega K.K., Tokyo, Japan, #GM2000)
- Falcon 96-well clear flat-bottom TC-treated culture microplates (Corning Inc., NY, USA, #353072)
- Nano-Glo HiBiT Extracellular Detection System (Promega K.K., Tokyo, Japan, #N2420)
- 50mg/mL Hygromycin B Solution (FUJIFILM Wako Pure Chemical Corporation, Osaka, Japan, #084-07681)
- HumanKine recombinant human IL-1 beta protein (Proteintech Japan, Tokyo, Japan, #HZ-1164)
- Nano-Glo Luciferase Assay System (Promega K.K., Tokyo, Japan, #N1130)

### Biological Resources

The sequences of template DNAs for in vitro transcription are presented in the supporting information. These sequences will also be available from DDBJ (Accession number: LC835990-LC836008). HuH-7 cells (RCB1366) were obtained from RIKEN BRC Cell Bank (Ibaraki, Japan). *Escherichia coli* DH5α (#9057) and HST08 (#9028) strains were obtained from Takara Bio (Shiga, Japan). pNL3.2.NF-κB-RE[NlucP/NF-κB-RE/Hygro] (#N1111) was obtained from Promega K.K. (Tokyo, Japan). pIR-CMVluc and pFerH-PBTP^19^ are kindly gifted from Prof. Yuriko Higuchi (Kyoto University).

### Cell culture

HuH-7 cells were maintained in D-MEM containing 10% fetal bovine serum and 0.5× penicillin–streptomycin. During passage, the cells were detached from the culture dishes by adding 0.25% trypsin/EDTA. The detached cells were then resuspended in the medium, and approximately 4 × 10^5^ cells were reseeded in new culture dishes.

### pDNA construction

PrimeSTAR Max DNA polymerase and KOD One PCR Master Mix were used to prepare inserts for PCR. The PCR and restriction digestion products were purified using the Monarch PCR & DNA Cleanup Kit. Cloning was performed using an In-Fusion HD Cloning Kit or In-Fusion Snap Assembly Master Mix. All pDNAs were amplified in *Escherichia coli* DH5α or HST08 using LB medium. Following amplification in *E. coli*, pDNAs were purified using the Monarch Plasmid Miniprep Kit or PureYield Plasmid Miniprep System. The concentration of purified pDNAs was measured using NanoDrop One. Sequences of the constructed pDNAs were analyzed via Sanger sequencing (Azenta Life Sciences, Tokyo, Japan).

### In vitro transcription of mRNAs

Template DNA for *in vitro* transcription was prepared via PCR using a procedure similar to that used for preparing inserts in pDNA construction. *In vitro* transcription was performed using the obtained template DNAs and either the MEGAscript T7 Transcription Kit or IVTpro mRNA Synthesis System. For all transcripts except for 1xMS2(U)site2-Luc2, the following components were used: 4.8 mM CleanCap Reagent AG (3’ OMe), 6 mM GTP, 6 mM ATP, 6 mM CTP, and 6 mM N^1^-methylpseudouridine-5’-triphosphate for *in vitro* transcription. For 1xMS2(U)site2-Luc2, 6 mM G(5’)ppp(5’)A RNA Cap Structure Analog, 1.5 mM GTP, 7.5 mM ATP, 7.5 mM CTP, and 7.5 mM N^1^-methylpseudouridine-5’-triphosphate were used. After transcription, TURBO DNase or DNase I was added to remove the template DNAs. The obtained mRNAs were purified using RNAClean XP and dephosphorylated using Quick CIP. Dephosphorylated mRNA was then purified using RNeasy Mini Kit or Monarch RNA Cleanup Kit. The concentration of purified mRNAs was measured using Qubit RNA Broad Range Assay Kit and Qubit 4 Fluorometer. The length of mRNAs was analyzed using an Agilent 2100 Bioanalyzer and Agilent RNA6000 Nano Kit.

### Dual-luciferase assay

A total of 5 × 10^3^ HuH-7 cells were seeded into CELLSTAR μClear 96-well TC F-bottom (chimney well) white microplates or Corning 96-well Flat Clear Bottom White Polystyrene TC-treated Microplates. One day after seeding, the cells were transfected with the mRNAs indicated in the figures and their legends using 0.3 μl/well of Lipofectamine MessengerMAX. For experiments on ligand-inducible translational regulation, the medium was replaced with a medium containing the indicated ligand ([Arg^8^]-vasopressin solution, prostaglandin E2, or bradykinin) approximately 4 h before transfection. One day after mRNA transfection, the luminescence of Luc2 and Nluc was measured using the Nano-Glo Dual-Luciferase Assay System and GloMax Navigator Microplate Luminometer. The quantity of transfected mRNAs in each experiment is indicated in the corresponding figure legends.

### Quantification of HiBiT-tagged secreted proteins

A total of 5 × 10^3^ HuH-7 cells were seeded into Falcon 96-well clear flat-bottom TC-treated culture microplates. A day after cell seeding, the medium was replaced with a medium containing prostaglandin E2. Approximately 3 h after medium replacement, cells were transfected with the mRNAs indicated in the figures and their legends using 0.3 μl/well of Lipofectamine MessengerMAX. The medium was collected at the indicated time points, and the wells were refilled with fresh media containing the indicated ligand concentrations. The HiBiT-tagged proteins in the collected medium were quantified using a Nano-Glo HiBiT Extracellular Detection System and GloMax Navigator Microplate Luminometer. The quantity of transfected mRNAs is presented in the figure legends.

### Quantification of NF-κB activity

A *piggyBac* transposon donor vector containing the NF-κB response element-driven *NlucP* gene, pIR-NFkBRE-NlucP, was constructed using pIR-CMVluc^19^ and pNL3.2.NF-κB-RE[NlucP/NF-κB-RE/Hygro] (Promega). HuH-7 cells were co-transfected with pIR-NFkBRE-NlucP and pFerH-PBTP^19^, a *piggyBac* transposase expression vector. These cells were then selected using hygromycin B to establish a stable cell line, HuH7-NFkBRE-NlucP. Subsequently, 5 × 10^3^ HuH7-NFkBRE-NlucP cells were seeded into CELLSTAR μClear 96-well TC F-bottom (chimney well) white microplates or Corning 96-well Flat Clear Bottom White Polystyrene TC-treated Microplates. The following day, the medium was replaced with a fresh medium containing prostaglandin E2 whose concentrations are indicated in the figures. Approximately 3 h after medium replacement, the cells were transfected with the indicated mRNAs using 0.3 μl/well of Lipofectamine MessengerMAX. After 24 h, the medium was replaced with fresh medium containing the indicated concentrations of prostaglandin E2 and human recombinant IL-1 beta protein. Two days post mRNA transfection, the luminescence of NlucP was measured using a Nano-Glo Luciferase Assay System and GloMax Navigator Microplate Luminometer. The quantity of transfected mRNAs in each experiment is indicated in the figure legends.

### Statistical Analyses

Statistical analyses were performed using Microsoft Excel and Python. For analysis using Python, pandas, SciPy, NumPy, statsmodels, Matplotlib, and seaborn libraries were utilized. For comparison between the two groups, the unpaired two-sided Student’s *t*-test was used. For comparison among three or more groups, Tukey’s multiple comparison test was used. To ensure reproducibility, three or more experiments were performed, and representative data are presented. All bar and line graphs indicate mean ± standard deviation (n = 4).

## Results

### Development of TEVp-sensitive translational regulator protein

To develop the system, we focused on G protein-coupled receptors (GPCRs), as they constitute the largest family of cell surface receptors with diverse target molecules^20,21^. Inspired by Tango and ChaCha, which are GPCR-based systems that regulate nuclear translocations^20–22^, we designed a translational regulation system composed of three mRNAs **(Figure 1)**. The first mRNA encodes a fusion protein of GPCR and TEVp. The second mRNA encodes a fusion protein of β-arrestin 2 (ARRB2), a TEVp-responsive degron (tevD)^6,7,23^, and caliciviral VPg-based translational activator (CaVT)^3,9,10,17,18^. CaVT is an artificial translational regulator that can be used for both translational repression and activation depending on the target mRNA design. While target mRNAs containing a 5′-cap and strong MS2-binding motif, such as 2xScMS2(C)-Luc2 in **Figure 2A**, are translationally repressed by CaVT, target mRNAs containing a translationally defective 5′-cap analog and weak MS2-binding motif, such as 1xMS2(U)site2-Luc2 in Figure 2A, are translationally activated. The third mRNA is a target for translational regulation and contains an MS2-binding motif and open reading frame (ORF) encoding a protein of interest. When a ligand is absent, CaVT binds to the target mRNA and represses or activates translation. In contrast, when a ligand is present, the ligand-GPCR binding induces the interaction between GPCR and ARRB2^20–22^, allowing TEVp to activate tevD. Activated tevD induces rapid degradation^6,7,23^ of CaVT, resulting in the release of target mRNA from CaVT.

**Figure 1.**
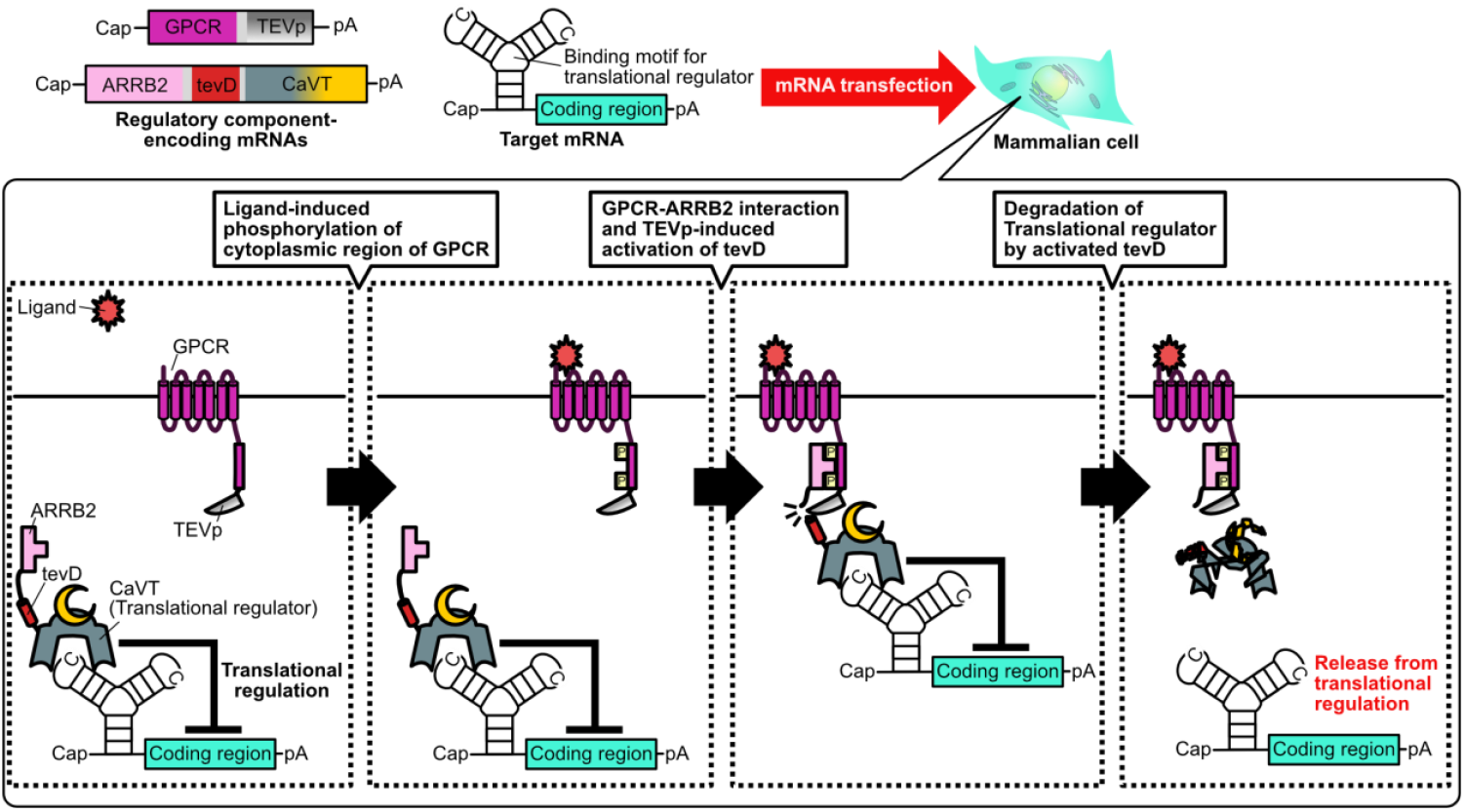
Schematic diagram of extracellular ligand-inducible translational regulation based on an engineered G protein-coupled receptor (GPCR) and a translational regulator protein. Before GPCR detects a ligand, the target mRNA is translationally repressed (or activated) by the ARRB2-fused translational regulator protein. Ligand binding to GPCR induces the phosphorylation of the cytoplasmic region of GPCR, followed by the interaction between the phosphorylated cytoplasmic region and ARRB2. This interaction allows TEVp to cleave the N-terminal cleavage site of tevD and induces tevD-mediated degradation of the translational regulator protein. Owing to this degradation, the target mRNA is released from the translational repression (or activation). TEVp: Tobacco Etch Virus protease; ARRB2: Beta-arrestin 2; tevD: TEV protease-responsive degron; CaVT: Caliciviral VPg-based Translational activator.

**Figure 2.**
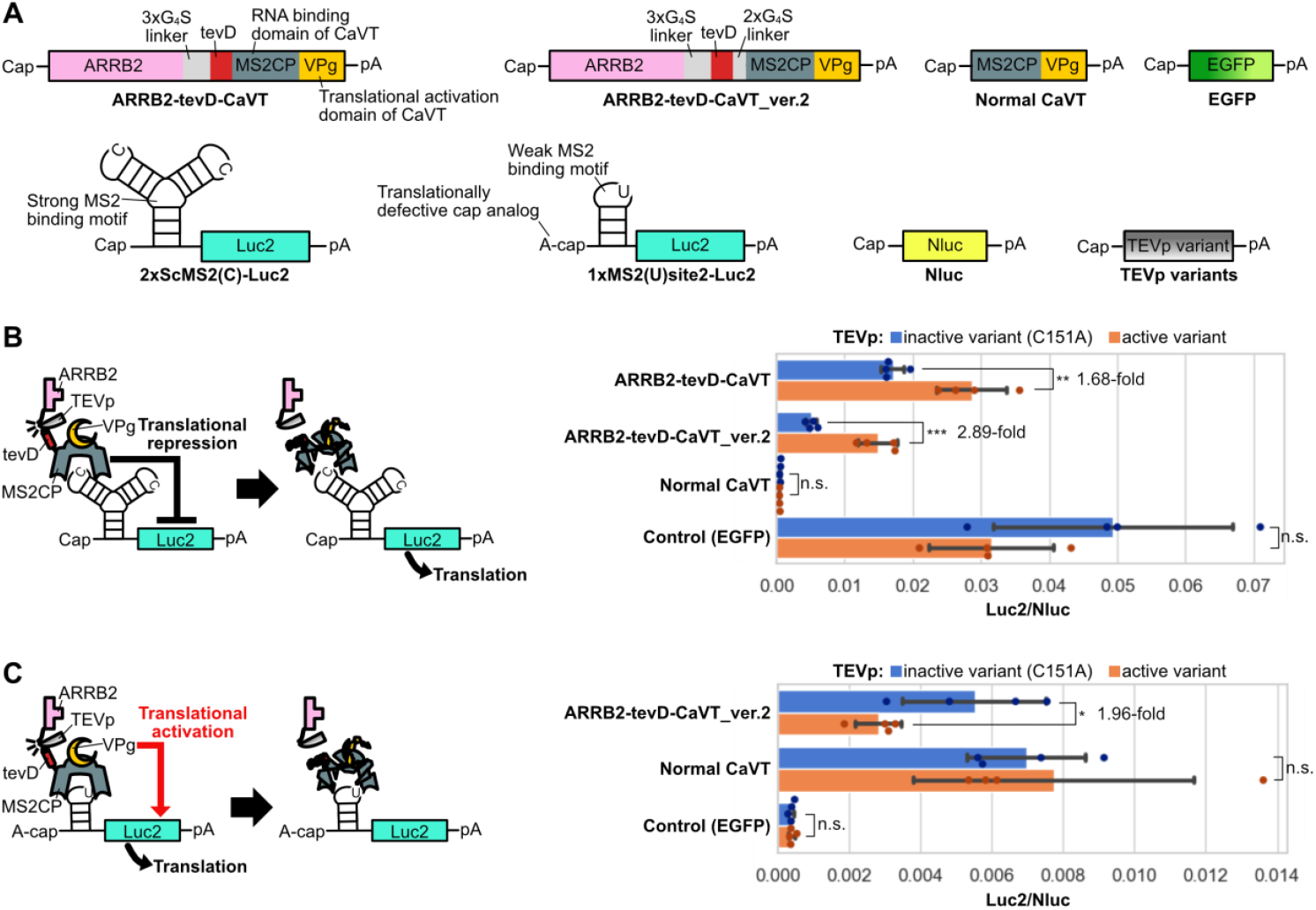
TEVp-responsiveness of ARRB2-tevD-fused CaVT. (A) Schematic diagram of mRNAs used to check the TEVp-responsiveness of CaVT fused with ARRB2-tevD. 1xMS2(U)site2-Luc2 mRNA was capped with A-cap, a translationally defective cap analog. All other mRNAs were capped with CleanCap AG (3’ OMe), a translationally active cap analog. (B) TEVp-responsive translational upregulation. HuH-7 cells in a 96-well plate were transfected with 2xScMS2(C)-Luc2 (20 ng/well), Nluc (1 ng/well), the indicated TEVp variant (10 ng/well), and the indicated CaVT variant or EGFP (70 ng/well) mRNAs. The bar graph represents the Luc2/Nluc ratio (mean ± SD, n = 4). ^**^, *P* < 0.01; ^***^, *P* < 0.001 by the unpaired two-sided Student’s *t*-test. (C) TEVp-responsive translational downregulation. The experiment followed the same procedure as (B) except that 1xMS2(U)site2-Luc2 mRNA was used instead of 2xScMS2(C)-Luc2 mRNA. The bar graph represents the Luc2/Nluc ratio (mean ± SD, n = 4). ^*^, *P* < 0.05 by the unpaired two-sided Student’s *t*-test. TEVp: Tobacco Etch Virus protease; ARRB2: Beta-arrestin 2; tevD: TEV protease-responsive degron; CaVT: Caliciviral VPg-based Translational activator.

To examine whether TEVp-induced degradation of tevD-fused CaVT could release the target mRNA from translational repression, we transfected HuH-7 cells with a target mRNA containing the strong MS2-binding motif (2xScMS2(C)-Luc2), ARRB2-tevD-CaVT, and TEVp or its C151A catalytically inactive mutant mRNAs^24^. As illustrated in **Figure 2B**, TEVp triggered an increase in the translation of 2xScMS2(C)-Luc2. However, the fold change was limited owing to the lower translational repression efficiency of ARRB2-tevD-CaVT compared to that of normal CaVT. This led us to suspect that the direct fusion of tevD and CaVT might negatively affect CaVT folding. To address this issue, we inserted a linker between tevD and CaVT to construct ARRB2-tevD-CaVT_ver.2. As this linker addition improved the fold change, we used ARRB2-tevD-CaVT_ver.2 in subsequent experiments.

As CaVT can be used for both translational repression and activation^3,9,10,17^, we sought to investigate if translational activation could be counteracted by TEVp. To this end, we transfected HuH-7 cells with a target mRNA containing a translationally defective cap analog and a weak MS2-binding motif (1xMS2(U)site2-Luc2), ARRB2-tevD-CaVT_ver.2, and TEVp mRNAs. Our results, depicted in **Figure 2C**, showed that TEVp cancelled translational activation by ARRB2-tevD-CaVT_ver.2.

### Development of TEVp-fused GPCRs for extracellular ligand-responsive translational regulation

To investigate extracellular ligand-responsive translational regulation, we fused TEVp to the C-terminus of AVP receptor 2 (AVPR2), which is a well-known GPCR **(Figure 3A)**. In addition to normal AVPR2-TEVp, we constructed three C-terminally truncated variants (AVPR2-TEVpΔ, AVPR2-TEVpΔT30A, and AVPR2-TEVpΔS135N). The C-terminal truncation of TEVp can decrease the affinity between TEVp and its target while still maintaining the cleavage activity^25^. Therefore, we hypothesized that truncation would improve fold change by decreasing the ligand-independent interaction between TEVp and tevD. Additionally, T30A and S135N mutations can increase the cleavage activity^25^. As depicted in **Figure 3B**, all AVPR2-TEVp variants exhibited AVP-responsive translational upregulation of 2xScMS2(C)-Luc2 when co-transfected with ARRB2-tevD-CaVT_ver.2. The AVPR2-TEVpΔT30A group displayed the highest fold change, prompting the selection of the T30A variant for subsequent experiments.

**Figure 3.**
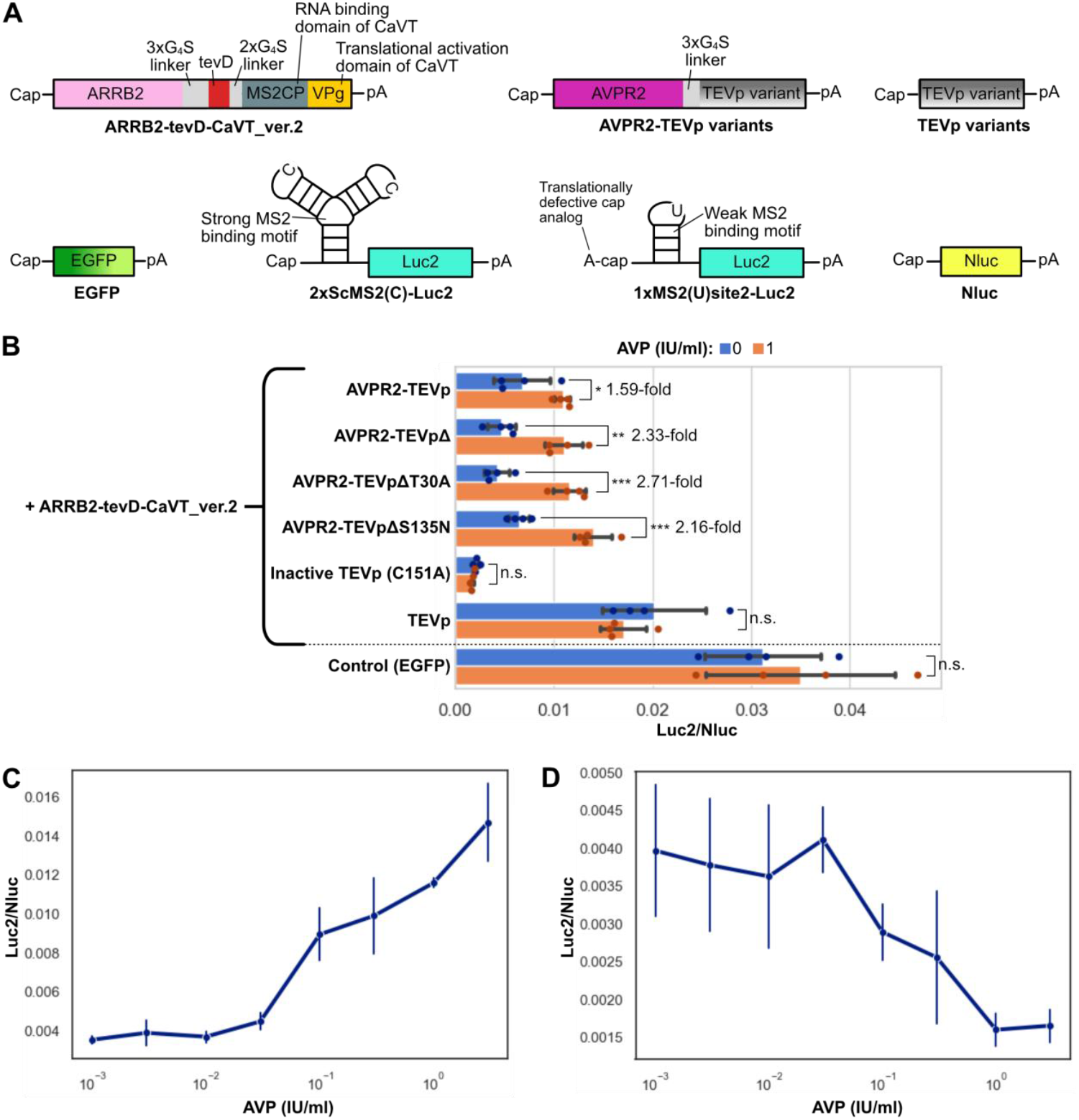
AVP-responsive translational upregulation and downregulation. (A) Schematic diagram of mRNAs utilized for AVP-responsive translational upregulation. (B) Comparison of different TEVp variants. HuH-7 cells were transfected with 2xScMS2(C)-Luc2 (20 ng/well), Nluc (1 ng/well), ARRB2-tevD-CaVT_ver.2 (70 ng/well), and the specified TEVp variant (10 ng/well) mRNAs in the presence or absence of AVP. The bar graph represents the Luc2/Nluc ratio (mean ± SD, n = 4). ^*^, *P* < 0.05; ^**^, *P* < 0.01; ^***^, *P* < 0.001 using the unpaired two-sided Student’s *t*-test. (C, D) Ligand concentration-dependency of AVP-responsive translational upregulation (C) and downregulation (D). HuH-7 cells were transfected with Nluc (1 ng/well), ARRB2-tevD-CaVT_ver.2 (70 ng/well), AVPR2-TEVpΔT30A (10 ng/well), and 2xScMS2(C)-Luc2 (20 ng/well) (C) or 1xMS2(U)site2-Luc2 (20 ng/well) (D) mRNAs. AVP: arginine vasopressin; AVPR2: arginine vasopressin receptor 2; TEVp: Tobacco Etch Virus protease; ARRB2: Beta-arrestin 2; tevD: TEV protease-responsive degron; CaVT: Caliciviral VPg-based Translational activator.

Next, we assessed the dependence of the system on ligand concentration. **Figure 3C** illustrates that the translation level increased in an AVP concentration-dependent manner. Additionally, when we used 1xMS2(U)site2-Luc2 instead of 2xScMS2(C)-Luc2, the translation level decreased in an AVP concentration-dependent manner (**Figure 3D**).

Next, we examined the versatility of the system. Given our focus on developing mRNA therapies for inflammatory diseases such as osteoarthritis^26,27^, we selected PGE2 and bradykinin (BK), which are known inflammatory markers^28^, as the ligands of interest. To this end, we substituted the receptor domain of the system from AVPR2 to PGE2 receptors (EP3 and EP4) and the BK receptor B2. As the cases of Tango and ChaCha^20–22^, we introduced the A343-S371 segment of AVPR2 to each receptor to enhance its interaction with ARRB2. Similar to the case of AVPR2, EP4 and B2 exhibited ligand-responsive translational upregulation (**Figure 4A and B**). In contrast, the isoform I of EP3 did not exhibit such upregulation (Figure 4A), although it reportedly interacts with ARRB2^29^. Given the importance of the C-terminal cytoplasmic region for GPCR–ARRB2 interaction, we substituted K351-R390 of EP3 with S327-S371 of AVPR2 to create an EP3-AVPR2 chimera (EP3V2). In contrast to EP3, EP3V2 enabled PGE2-responsive translational upregulation (**Figure 4C**).

**Figure 4.**
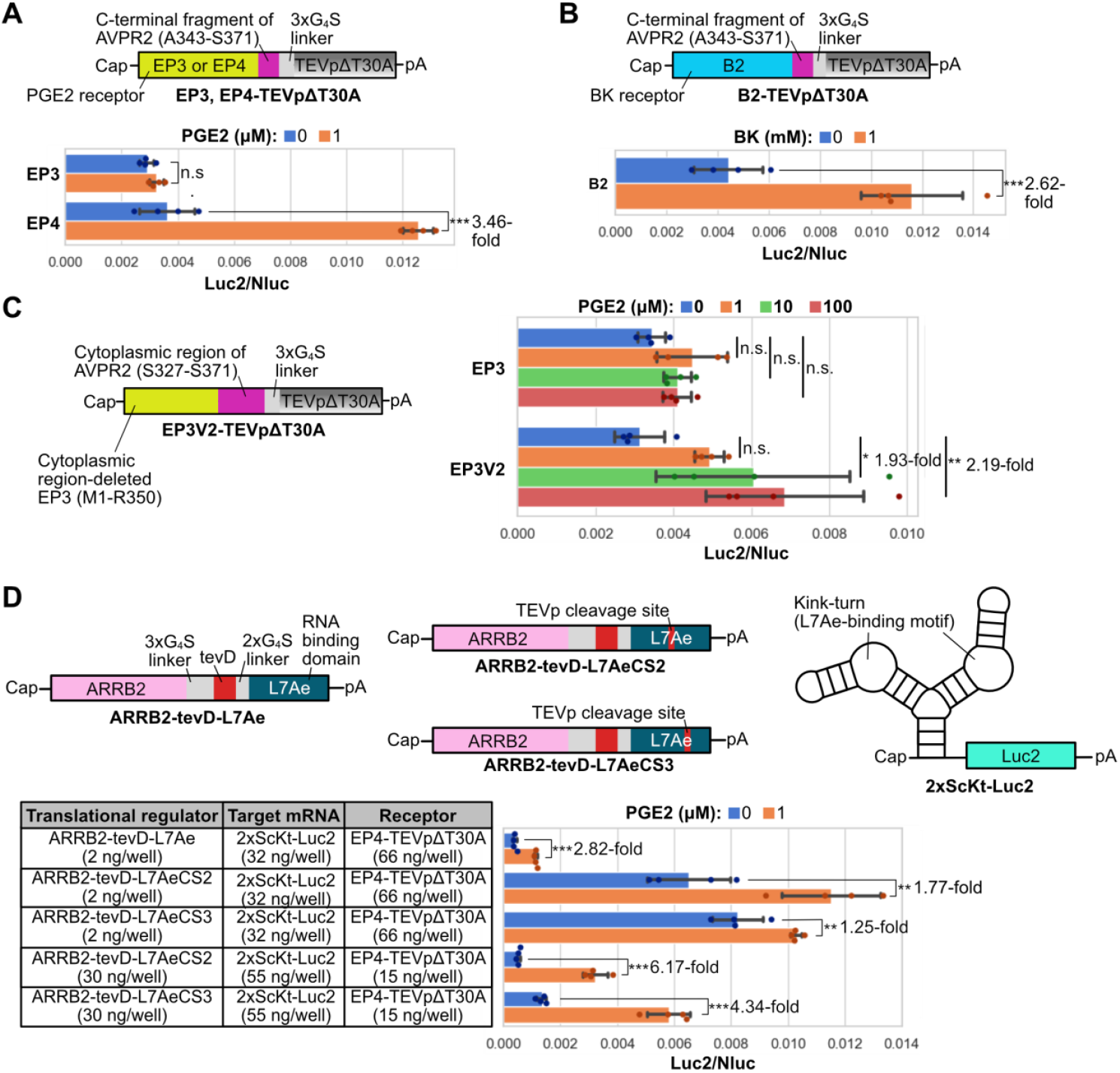
Translational upregulation utilizing GPCRs for inflammatory marker detection. (A, B, C) Inflammatory marker-responsive translational upregulation using ARRB2-tevD-CaVT_ver.2 and engineered receptors. PGE2 receptors (EP3 or EP4) (A), BK receptor B2 (B), or EP3-AVPR2 chimeric receptor (EP3V2) (C) were fused with TEVpΔT30A. The experiments were conducted following the same procedure as in Figure 3C, except for the ligands and their receptors. (D) PGE2-responsive translational upregulation using ARRB2-tevD-L7Ae and its variants. HuH-7 cells were transfected with Nluc (1 ng/well), indicated ARRB2-tevD-L7Ae variants (2 or 30 ng/well), EP4-TEVpΔT30A (66 or 15 ng/well), and 2xScKt-Luc2 (32 or 55 ng/well) mRNAs. The bar graph represents the Luc2/Nluc ratio (mean ± SD, n = 4). ^***^, *P* < 0.001 using the unpaired two-sided Student’s *t*-test (A, B, D). ^*^, *P* < 0.05; ^**^, *P* < 0.01 using Tukey’s multiple comparison test (C). AVPR2: arginine vasopressin receptor 2; TEVp: Tobacco Etch Virus protease; PGE2: Prostaglandin E2; BK: Bradykinin.

Furthermore, we also investigated whether this regulatory mechanism can be applied to a translational regulator protein other than CaVT. For this purpose, we substituted the translational regulator domain of ARRB2-tevD-CaVT_ver.2 from CaVT to L7Ae, an RNA-binding protein that represses the translation of kink-turn motif-embedded mRNAs^30^. This new translational regulator protein, named ARRB2-tevD-L7Ae, showed high translational repression efficiency in the absence of the ligand, but its ligand responsiveness was relatively lower than that of ARRB2-tevD-CaVT_ver.2 even after optimizing the transfection ratio of ARRB2-tevD-L7Ae and the engineered receptor (**Supplementary Figure S1**). Therefore, we added a TEVp-cleavage site after amino acid residue P56 or K77 of L7Ae^31^ to obtain ARRB2-tevD-L7AeCS2 and CS3, respectively. Although the insertion of the TEVp-cleavage site decreased the translational repression efficiency of L7Ae, after optimizing the transfection ratio, ARRB2-tevD-L7AeCS2 and CS3 showed higher fold change than ARRB2-tevD-L7Ae (**Figure 4D**).

### Suppression of NF-κB activation by a translationally regulated anti-inflammatory protein

After previously discovering that inflammatory diseases can be alleviated by an interleukin (IL)-1 receptor antagonist (IL-1Ra)-encoding mRNA^26^, we hypothesized that increasing IL-1Ra production upon detecting the exacerbation of inflammatory diseases could present a promising treatment approach. To test this hypothesis, we transfected HuH-7 cells with EP4-TEVpΔT30A, ARRB2-tevD-CaVT_ver.2, and an mRNA containing a strong MS2-binding motif and a HiBiT-tagged *IL-1Ra* ORF (2xScMS2(C)-IL1Ra-HiBiT) **(Figure 5A). Figure 5B** illustrates that PGE2 increased IL-1Ra secretion for at least three days. In contrast, IL1Ra-HiBiT mRNA lacking the MS2-binding motif did not exhibit PGE2-induced upregulation (**Supplementary Figure S2**).

**Figure 5.**
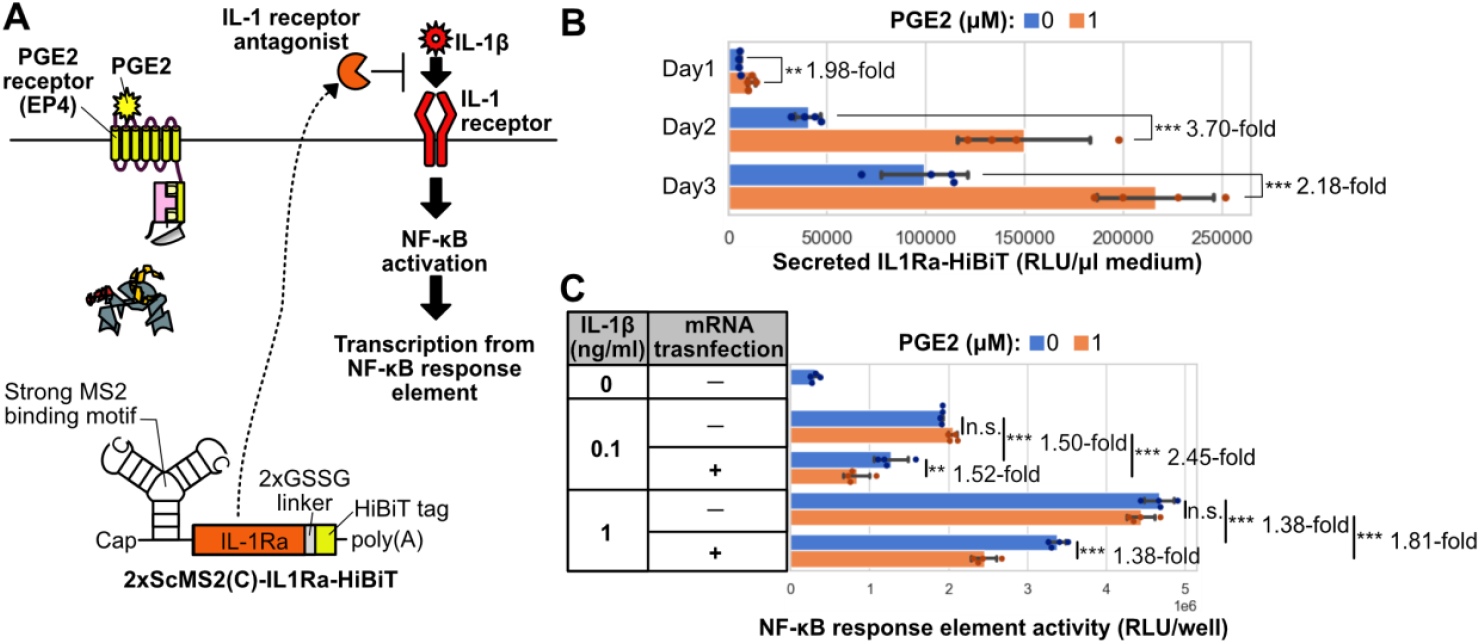
PGE2-responsive translational upregulation of an anti-inflammatory protein. (A) Schematic diagram of NF-κB activity repression by the translationally activated IL-1Ra. (B) IL1Ra-HiBiT secretion from HuH-7 cells transfected with 2xScMS2(C)-IL1Ra-HiBiT (20 ng/well), ARRB2-tevD-CaVT_ver.2 (70 ng/well), and EP4-TEVpΔT30A (10 ng/well). The bar graph represents HiBiT luminescence in the medium (mean ± SD, n = 4). **, *P* < 0.01; ***, *P* < 0.001 using the unpaired two-sided Student’s *t*-test. (C) NF-κB activity analysis using the NF-κB reporter cells. The reporter cells were transfected with the same mRNA cocktail, as shown in (B), and treated with IL-1β. The bar graph represents NlucP luminescence (mean ± SD, n = 4). **, *P* < 0.01; ***, *P* < 0.001 using Tukey’s multiple comparison test. IL: Interleukin; IL-1Ra: IL-1 receptor antagonist; PGE2: Prostaglandin E2.

Subsequently, we investigated whether the translational upregulation of IL-1Ra can suppress IL-1β, a proinflammatory cytokine. As IL-1β can activate NF-κB in HuH-7 cells^32^, we inserted NF-κB response element-driven *NlucP* genes into the genome of HuH-7 cells to monitor NF-κB activation (Figure 5A). The established stable cell line (HuH7-NFkBRE-NlucP) was transfected with 2xScMS2(C)-IL1Ra-HiBiT, ARRB2-tevD-CaVT_ver.2, and EP4-TEVpΔT30A mRNAs in the absence or presence of PGE2. In the presence of PGE2, a 2.45-fold decrease in NlucP expression was observed in the mRNA-transfected group, compared to the untransfected group, when NF-κB activation was induced by 0.1 ng/ml IL-1β. This decrease in NlucP expression was also evident when the concentration of IL-1β increased to 1 ng/ml; however, the fold change decreased to 1.81 (**Figure 5C**).

## Discussion

While the current major applications of synthetic mRNAs are vaccines to express antigens for infectious diseases and cancers, mRNA drugs aimed to express proteins also hold promise.^1^ However, compared to mRNA vaccines, therapeutic windows are more important in the context of mRNA drugs. Specifically, although protein production levels below the therapeutic window result in insufficient therapeutic effects, excessive protein production may cause adverse effects. Therefore, a system capable of producing appropriate amounts of therapeutic proteins according to disease states is required. Given the importance of extracellular biomolecules, such as hormones and lipid mediators, as markers indicating disease states, in the present study, we developed an extracellular ligand-responsive translational regulation system utilizing GPCRs. As GPCRs constitute the largest family of cell surface receptors, the range of disease markers detectable by GPCRs is considerably large ^20,21^. Although the data obtained with EP3-TEVpΔT30A (Figure 4A) indicated that not all GPCRs are suitable as components of our translational regulation system, the engineering of the C-terminal cytoplasmic region of EP3 enabled EP3-mediated translational regulation (Figure 4C). This result suggests that even if a GPCR for a ligand to be detected is incompatible with our system, appropriate engineering can enable its utilization.

The major applications of our system are mRNA drugs to express therapeutic proteins. However, it can also be used to improve mRNA vaccines. One possible application is improving the safety of mRNA vaccines by adjusting inflammation levels. Inflammation induced by mRNA vaccines is related to both immunogenicity and adverse effects, and appropriate inflammation levels are important for successful immunization without severe adverse effects^33^. However, these inflammation levels vary among vaccine recipients. We anticipate that the inflammatory marker-responsive production of anti-inflammatory proteins will help adjust inflammation levels in each recipient.

Like other translational regulation systems^8,9,12,15,16^, one limitation of our system is its leaky expression (Figure 5B). As ARRB2-tevD-CaVT_ver.2 and target mRNAs were transfected concurrently, translational initiation of both mRNAs should theoretically occur simultaneously. Thus, even in the absence of a ligand, target mRNA translation can be repressed only after the production of a sufficient amount of ARRB2-tevD-CaVT_ver.2. To avoid this issue, future engineering aimed at delaying the translational initiation of target mRNAs is desirable.

In our system, we used CaVT and L7Ae as translational regulator proteins. However, other RNA-binding proteins and translational modulator proteins can also be used as components of our system. For example, LIN28A^16^, and PP7 coat protein^34^ can be used as an RNA-binding protein instead of MS2CP. Similarly, murine norovirus VPg^35^ and rotavirus NSP3^5^ can be used as a positive translational modulator instead of feline calicivirus VPg. Additionally, some proteins, such as CNOT7^11^, DDX6^4^, and Scd6^35^ can be used as a negative translational modulator. If it is necessary to regulate multiple target mRNAs encoding different therapeutic proteins, using multiple combinations of translational regulators and their target RNA motifs is a promising approach. Such an approach will allow separate regulations of multiple therapeutic protein production by a single ligand.

In the present study, we used conventional synthetic mRNAs with relatively short half-lives. However, there are several novel mRNA types that enable extended protein expression durations, such as self-amplifying mRNAs^36^, poly(A) tail-modified mRNAs^37,38^, and circular RNAs^39^. For these long-lasting mRNAs, disease marker-responsive translational regulation is more important, as the possibility of disease state changes during protein expression duration is higher. Given that the translational regulation of self-amplifying mRNAs and circular RNAs by RNA-binding proteins has been already reported^4,8,11,40^, we believe that our system can also be integrated with these long-lasting RNAs.

In summary, we developed an extracellular ligand-responsive translational regulation system that can be used for both upregulation and downregulation. The target ligand can be changed by substituting the GPCR domain, making this system versatile and adaptable for various disease markers. We expect this system to pave the way for next-generation mRNA drugs, enabling autonomous optimization of therapeutic effects.

## Acknowledgements

We thank Prof. Yuriko Higuchi (Kyoto University) for the *piggyBac* transposon vectors (pIR-CMVluc and pFerH-PBTP). We also thank Ms. Yoko Hasegawa (TMDU) for her technical support. Some reagents were provided as prizes by Nippon Genetics. This work was supported by the Japan Society for the Promotion of Science (JSPS) KAKENHI [JP19K20696 and JP23K11843 to H.N., JP23K28426 to K.I.], the Japan Agency for Medical Research and Development (AMED) [24fk0310515s0503 and JP223fa627002 to K.I.], and Tokyo Medical and Dental University (TMDU) IBB Medical & Dental Research Grant. This research was also supported in part by Axcelead, Inc.

## Data Availability Statement

Template DNA sequences of mRNAs used in this study are presented in the supporting information. These sequences will also be available from DDBJ (Accession number: LC835990-LC836008). If additional data are required, they would be available from the corresponding authors upon reasonable request.

## References

1. Parhiz, H., Atochina-Vasserman, E. N. & Weissman, D. mRNA-based therapeutics: looking beyond COVID-19 vaccines. Lancet 403, 1192–1204 (2024).

2. Matsui, A., Uchida, S., Ishii, T., Itaka, K. & Kataoka, K. Messenger RNA-based therapeutics for the treatment of apoptosis-associated diseases. Sci. Rep. 5, 15810 (2015).

3. Nakanishi, H. & Saito, H. Caliciviral protein-based artificial translational activator for mammalian gene circuits with RNA-only delivery. Nat. Commun. 11, 1297 (2020).

4. Wagner, T. E., Becraft, J. R., Bodner, K., Teague, B., Zhang, X., Woo, A., Porter, E., Alburquerque, B., Dobosh, B., Andries, O., Sanders, N. N., Beal, J., Densmore, D., Kitada, T. & Weiss, R. Small-molecule-based regulation of RNA-delivered circuits in mammalian cells. Nat. Chem. Biol. 14, 1043–1050 (2018).

5. Shao, J., Li, S., Qiu, X., Jiang, J., Zhang, L., Wang, P., Si, Y., Wu, Y., He, M., Xiong, Q., Zhao, L., Li, Y., Fan, Y., Viviani, M., Fu, Y., Wu, C., Gao, T., Zhu, L., Fussenegger, M., Wang, H. & Xie, M. Engineered poly(A)-surrogates for translational regulation and therapeutic biocomputation in mammalian cells. Cell Res. 34, 31–46 (2024).

6. Yang, J. & Ding, S. Engineering L7Ae for RNA-Only Delivery Kill Switch Targeting CMS2 Type Colorectal Cancer Cells. ACS Synth. Biol. 10, 1095–1105 (2021).

7. Yang, J. & Ding, S. Chimeric RNA-binding protein-based killing switch targeting hepatocellular carcinoma cells. Mol. Ther. Nucleic Acids 25, 683–695 (2021).

8. Mc Cafferty, S., De Temmerman, J., Kitada, T., Becraft, J. R., Weiss, R., Irvine, D. J., Devreese, M., De Baere, S., Combes, F. & Sanders, N. N. In Vivo Validation of a Reversible Small Molecule-Based Switch for Synthetic Self-Amplifying mRNA Regulation. Mol. Ther. 29, 1164–1173 (2021).

9. Nakanishi, H., Yoshii, T., Kawasaki, S., Hayashi, K., Tsutsui, K., Oki, C., Tsukiji, S. & Saito, H. Light-controllable RNA-protein devices for translational regulation of synthetic mRNAs in mammalian cells. Cell Chem. Biol. 28, 662-674.e5 (2021).

10. Nakanishi, H., Yoshii, T., Tsukiji, S. & Saito, H. A protocol to construct RNA-protein devices for photochemical translational regulation of synthetic mRNAs in mammalian cells. STAR Protoc. 3, 101451 (2022).

11. Wroblewska, L., Kitada, T., Endo, K., Siciliano, V., Stillo, B., Saito, H. & Weiss, R. Mammalian synthetic circuits with RNA binding proteins for RNA-only delivery. Nat. Biotechnol. 33, 839–841 (2015).

12. Fujita, Y., Hirosawa, M., Hayashi, K., Hatani, T., Yoshida, Y., Yamamoto, T. & Saito, H. A versatile and robust cell purification system with an RNA-only circuit composed of microRNA-responsive ON and OFF switches. Sci. Adv. 8, eabj1793 (2022).

13. Zhao, E. M., Mao, A. S., de Puig, H., Zhang, K., Tippens, N. D., Tan, X., Ran, F. A., Han,, Nguyen, P. Q., Chory, E. J., Hua, T. Y., Ramesh, P., Thompson, D. B., Oh, C. Y., Zigon, E. S., English, M. A. & Collins, J. J. RNA-responsive elements for eukaryotic translational control. Nat. Biotechnol. 40, 539–545 (2022).

14. Jiang, K., Koob, J., Chen, X. D., Krajeski, R. N., Zhang, Y., Volf, V., Zhou, W., Sgrizzi, S. R., Villiger, L., Gootenberg, J. S., Chen, F. & Abudayyeh, O. O. Programmable eukaryotic protein synthesis with RNA sensors by harnessing ADAR. Nat Biotechnol 41, 698–707 (2023).

15. Ning, H., Liu, G., Li, L., Liu, Q., Huang, H. & Xie, Z. Rational design of microRNA-responsive switch for programmable translational control in mammalian cells. Nat. Commun. 14, 7193 (2023).

16. Kawasaki, S., Fujita, Y., Nagaike, T., Tomita, K. & Saito, H. Synthetic mRNA devices that detect endogenous proteins and distinguish mammalian cells. Nucleic Acids Res. 45, e117–e117 (2017).

17. Nakanishi, H., Saito, H. & Itaka, K. Versatile Design of Intracellular Protein-Responsive Translational Regulation System for Synthetic mRNA. ACS Synth. Biol. 11, 1077–1085 (2022).

18. Yang, T., Nakanishi, H. & Itaka, K. Development of a new caged intein for multi-input conditional translation of synthetic mRNA. Sci. Rep. 14, 9988 (2024).

19. Nakanishi, H., Higuchi, Y., Kawakami, S., Yamashita, F. & Hashida, M. piggyBac Transposon-mediated Long-term Gene Expression in Mice. Mol. Ther. 18, 707–714 (2010).

20. Kroeze, W. K., Sassano, M. F., Huang, X.-P., Lansu, K., McCorvy, J. D., Giguère, P. M., Sciaky, N. & Roth, B. L. PRESTO-Tango as an open-source resource for interrogation of the druggable human GPCRome. Nat. Struct. Mol. Biol. 22, 362–369 (2015).

21. Kipniss, N. H., Dingal, P. C. D. P., Abbott, T. R., Gao, Y., Wang, H., Dominguez, A. A., Labanieh, L. & Qi, L. S. Engineering cell sensing and responses using a GPCR-coupled CRISPR-Cas system. Nat. Commun. 8, 2212 (2017).

22. Barnea, G., Strapps, W., Herrada, G., Berman, Y., Ong, J., Kloss, B., Axel, R. & Lee, K. J. The genetic design of signaling cascades to record receptor activation. Proc. Natl. Acad. Sci. U. S. A. 105, 64–69 (2008).

23. Taxis, C., Stier, G., Spadaccini, R. & Knop, M. Efficient protein depletion by genetically controlled deprotection of a dormant N-degron. Mol. Syst. Biol. 5, 267 (2009).

24. Phan, J., Zdanov, A., Evdokimov, A. G., Tropea, J. E., Peters, H. K., Kapust, R. B., Li, M., Wlodawer, A. & Waugh, D. S. Structural Basis for the Substrate Specificity of Tobacco Etch Virus Protease. J. Biol. Chem. 277, 50564–50572 (2002).

25. Sanchez, M. I. & Ting, A. Y. Directed evolution improves the catalytic efficiency of TEV protease. Nat. Methods 17, 167–174 (2020).

26. Deng, J., Fukushima, Y., Nozaki, K., Nakanishi, H., Yada, E., Terai, Y., Fueki, K. & Itaka, K. Anti-Inflammatory Therapy for Temporomandibular Joint Osteoarthritis Using mRNA Medicine Encoding Interleukin-1 Receptor Antagonist. Pharmaceutics 14, 1785 (2022).

27. Aini, H., Itaka, K., Fujisawa, A., Uchida, H., Uchida, S., Fukushima, S., Kataoka, K., Saito, T., Chung, U. & Ohba, S. Messenger RNA delivery of a cartilage-anabolic transcription factor as a disease-modifying strategy for osteoarthritis treatment. Sci. Rep. 6, 18743 (2016).

28. Wieland, H. A., Michaelis, M., Kirschbaum, B. J. & Rudolphi, K. A. Osteoarthritis — an untreatable disease? Nat. Rev. Drug Discov. 4, 331–344 (2005).

29. Bilson, H. A., Mitchell, D. L. & Ashby, B. Human prostaglandin EP3 receptor isoforms show different agonist-induced internalization patterns. FEBS Lett. 572, 271–275 (2004).

30. Saito, H., Kobayashi, T., Hara, T., Fujita, Y., Hayashi, K., Furushima, R. & Inoue, T. Synthetic translational regulation by an L7Ae–kink-turn RNP switch. Nat. Chem. Biol. 6, 71–78 (2010).

31. Cella, F., Wroblewska, L., Weiss, R. & Siciliano, V. Engineering protein-protein devices for multilayered regulation of mRNA translation using orthogonal proteases in mammalian cells. Nat. Commun. 9, 4392 (2018).

32. Zhang, T., Guo, C.-J., Li, Y., Douglas, S. D., Qi, X.-X., Song, L. & Ho, W.-Z. Interleukin-1β induces macrophage inflammatory protein-1β expression in human hepatocytes. Cell Immunol. 226, 45–53 (2003).

33. Lee, J., Woodruff, M. C., Kim, E. H. & Nam, J.-H. Knife’s edge: Balancing immunogenicity and reactogenicity in mRNA vaccines. Exp. Mol. Med. 55, 1305–1313 (2023).

34. Ono, H., Kawasaki, S. & Saito, H. Orthogonal Protein-Responsive mRNA Switches for Mammalian Synthetic Biology. ACS Synth. Biol. 9, 169–174 (2020).

35. Tan, K., Hu, Y., Liang, Z., Li, C. Y., Yau, W. L. & Kuang, Y. Dual Input-Controlled Synthetic mRNA Circuit for Bidirectional Protein Expression Regulation. ACS Synth. Biol. 12, 2516–2523 (2023).

36. Huysmans, H., Zhong, Z., De Temmerman, J., Mui, B. L., Tam, Y. K., Mc Cafferty, S., Gitsels, A., Vanrompay, D. & Sanders, N. N. Expression Kinetics and Innate Immune Response after Electroporation and LNP-Mediated Delivery of a Self-Amplifying mRNA in the Skin. Mol. Ther. Nucleic Acids 17, 867–878 (2019).

37. Li, C. Y., Liang, Z., Hu, Y., Zhang, H., Setiasabda, K. D., Li, J., Ma, S., Xia, X. & Kuang, Y. Cytidine-containing tails robustly enhance and prolong protein production of synthetic mRNA in cell and in vivo. Mol. Ther. Nucleic Acids 30, 300–310 (2022).

38. Chen, H., Liu, D., Guo, J., Aditham, A., Zhou, Y., Tian, J., Luo, S., Ren, J., Hsu, A., Huang, J., Kostas, F., Wu, M., Liu, D. R. & Wang, X. Branched chemically modified poly(A) tails enhance the translation capacity of mRNA. Nat. Biotechnol. 1–10 (2024). doi:10.1038/s41587-024-02174-7

39. Wesselhoeft, R. A., Kowalski, P. S. & Anderson, D. G. Engineering circular RNA for potent and stable translation in eukaryotic cells. Nat. Commun. 9, 2629 (2018).

40. Kameda, S., Ohno, H. & Saito, H. Synthetic circular RNA switches and circuits that control protein expression in mammalian cells. Nucleic Acids Res. 51, e24 (2023).

