## Supplementary Figures and Methods for "Extracellular Ligand-Responsive Translational Regulation of Synthetic mRNAs using Engineered Receptors"

- Supplementary Figure S1 | Comparison of ARRB2-tevD-CaVT_ver.2 and ARRB2-tevD-L7Ae in AVP-responsive translational upregulation.
- Supplementary Figure S2 | Time course of IL-1Ra secretion from cells transfected with the mRNA lacking an MS2-binding motif.

**Supplementary Materials and Methods**

- Full sequences of template DNAs for in vitro transcription.

**Supplementary Figures**

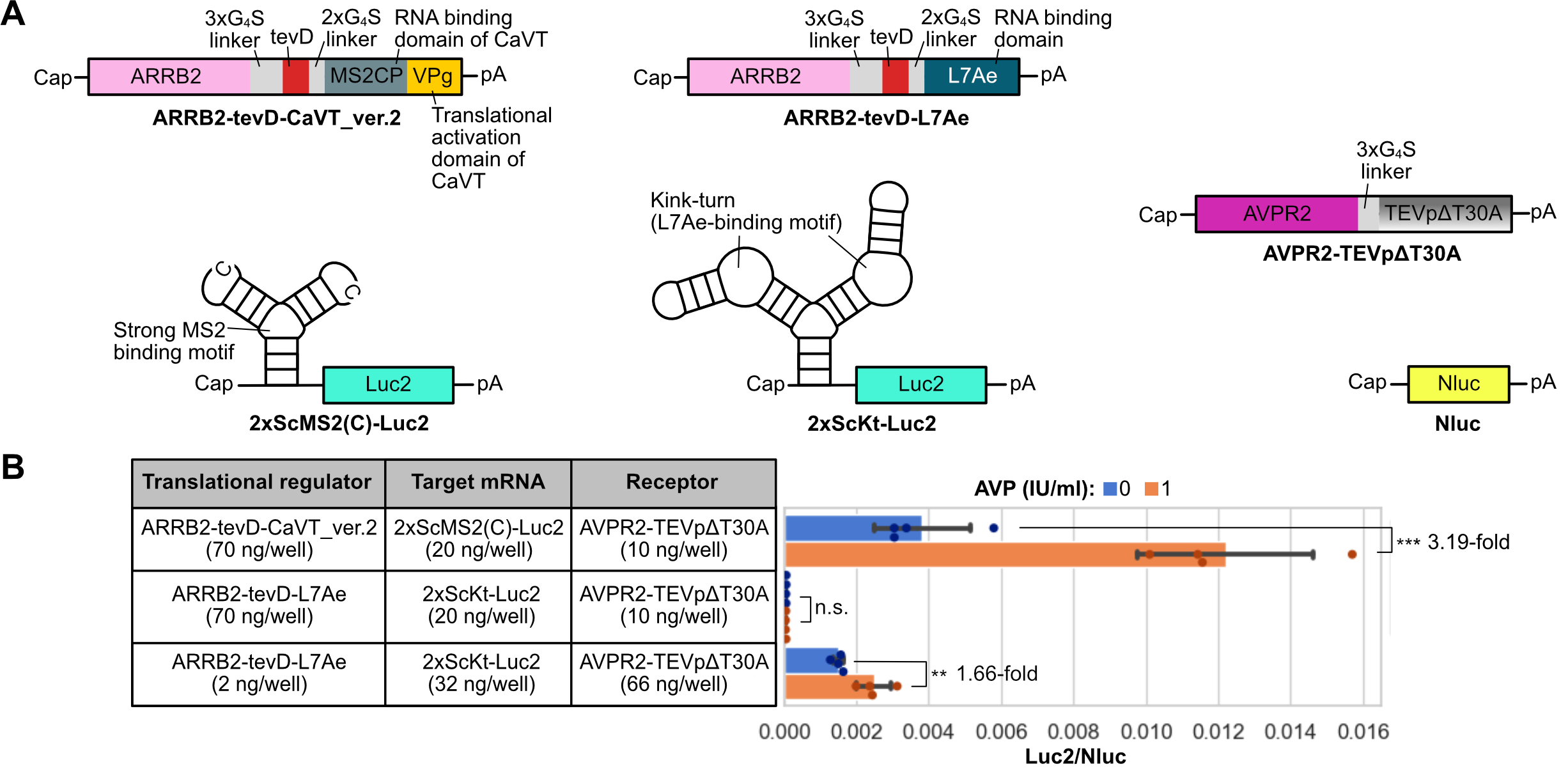

**Supplementary Figure S1 | Comparison of ARRB2-tevD-CaVT_ver.2 and ARRB2-tevD-L7Ae in ligand-responsive translational upregulation.**

(A) Schematic diagram of mRNAs to compare ARRB2-tevD-CaVT_ver.2 and ARRB2-tevD-L7Ae in AVP-responsive translational upregulation. (B) HuH-7cells were transfected with Nluc (1 ng/well), indicated translational regulator (70 or 2 ng/well), target (20 or 32 ng/well), and receptor (10 or 66 ng/well) mRNAs in the presence or absence of AVP. The bar graph represents the Luc2/Nluc ratio (mean ± SD, n = 4). **, *P* < 0.01; ***, *P* < 0.001 using the unpaired two-sided Student’s *t*-test.

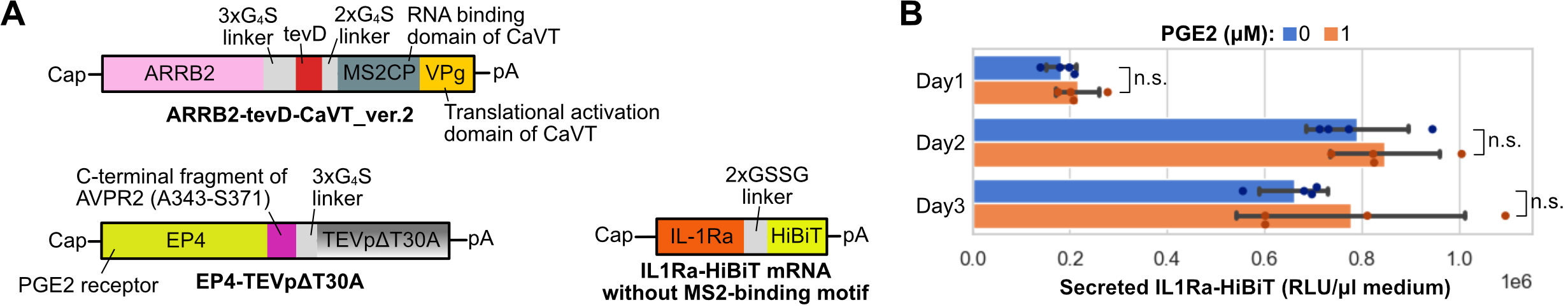

**Supplementary Figure S2 | Time course of IL-1Ra secretion from cells transfected with the mRNA lacking an MS2-binding motif.**

(A) Schematic diagram of mRNAs to check IL1Ra-HiBiT translation from a conventional mRNA. (B) Time course of IL1Ra-HiBiT secretion. HuH-7 cells were seeded onto a 96-well plate. One day after cell seeding, the medium was replaced with medium containing 0 or 1 μM of PGE2. After the medium replacement, cells were transfected with IL1Ra-HiBiT (20 ng/well), Nluc (1 ng/well), ARRB2-tevD-CaVT_ver.2 (70 ng/well), and EP4-TEVpΔT30A (10 ng/well) mRNAs. The medium was collected at the indicated time points. The bar graph shows HiBiT luminescence in 50 μl of medium samples (mean ± SD, n = 4).

**Supplementary Materials and Methods**

**Full sequences of template DNAs for in vitro transcription**

- **2xScMS2(C)-Luc2**

T7 promoter (for CleanCap AG Reagent): 11-30

Stabilized 2xMS2-binding motif (C-variant): 51-117

Kozak sequence (including start codon): 125-133

Luc2: 134-1780

Stop codon: 1781-1783

[CAGTGAATTGTAATACGACTCACTATAAGGTCAGATCCGCTAGCGGATCCGGGAGCAGGTGAGGATCACCCATCTGCCACGAGCGAGGTGAGGATCACCCATCTCGCTCGTGTTCCCACCGGTCGCCACCATGGAAGATGCCAAAAACATTAAGAAGGGCCCAGCGCCATTCTACCCACTCGAAGACGGGACCGCCGGCGAGCAGCTGCACAAAGCCATGAAGCGCTACGCCCTGGTGCCCGGCACCATCGCCTTTACCGACGCACATATCGAGGTGGACATTACCTACGCCGAGTACTTCGAGATGAGCGTTCGGCTGGCAGAAGCTATGAAGCGCTATGGGCTGAATACAAACCATCGGATCGTGGTGTGCAGCGAGAATAGCTTGCAGTTCTTCATGCCCGTGTTGGGTGCCCTGTTCATCGGTGTGGCTGTGGCCCCAGCTAACGACATCTACAACGAGCGCGAGCTGCTGAACAGCATGGGCATCAGCCAGCCCACCGTCGTATTCGTGAGCAAGAAAGGGCTGCAAAAGATCCTCAACGTGCAAAAGAAGCTACCGATCATACAAAAGATCATCATCATGGATAGCAAGACCGACTACCAGGGCTTCCAAAGCATGTACACCTTCGTGACTTCCCATTTGCCACCCGGCTTCAACGAGTACGACTTCGTGCCCGAGAGCTTCGACCGGGACAAAACCATCGCCCTGATCATGAACAGTAGTGGCAGTACCGGATTGCCCAAGGGCGTAGCCCTACCGCACCGCACCGCTTGTGTCCGATTCAGTCATGCCCGCGACCCCATCTTCGGCAACCAGATCATCCCCGACACCGCTATCCTCAGCGTGGTGCCATTTCACCACGGCTTCGGCATGTTCACCACGCTGGGCTACTTGATCTGCGGCTTTCGGGTCGTGCTCATGTACCGCTTCGAGGAGGAGCTATTCTTGCGCAGCTTGCAAGACTATAAGATTCAATCTGCCCTGCTGGTGCCCACACTATTTAGCTTCTTCGCTAAGAGCACTCTCATCGACAAGTACGACCTAAGCAACTTGCACGAGATCGCCAGCGGCGGGGCGCCGCTCAGCAAGGAGGTAGGTGAGGCCGTGGCCAAACGCTTCCACCTACCAGGCATCCGCCAGGGCTACGGCCTGACAGAAACAACCAGCGCCATTCTGATCACCCCCGAAGGGGACGACAAGCCTGGCGCAGTAGGCAAGGTGGTGCCCTTCTTCGAGGCTAAGGTGGTGGACTTGGACACCGGTAAGACACTGGGTGTGAACCAGCGCGGCGAGCTGTGCGTCCGTGGCCCCATGATCATGAGCGGCTACGTTAACAACCCCGAGGCTACAAACGCTCTCATCGACAAGGACGGCTGGCTGCACAGCGGCGACATCGCCTACTGGGACGAGGACGAGCACTTCTTCATCGTGGACCGGCTGAAGAGCCTGATCAAATACAAGGGCTACCAGGTAGCCCCAGCCGAACTGGAGAGCATCCTGCTGCAACACCCCAACATCTTCGACGCCGGGGTCGCCGGCCTGCCCGACGACGATGCCGGCGAGCTGCCCGCCGCAGTCGTCGTGCTGGAACACGGTAAAACCATGACCGAGAAGGAGATCGTGGACTATGTGGCCAGCCAGGTTACAACCGCCAAGAAGCTGCGCGGTGGTGTTGTGTTCGTGGACGAGGTGCCTAAAGGACTGACCGGCAAGTTGGACGCCCGCAAGATCCGCGAGATTCTCATTAAGGCCAAGAAGGGCGGCAAGATCGCCGTGTAAATCTAGACCTTCTGCGGGGCTTGCCTTCTGGCCATGCCCTTCTTCTCTCCCTTGCACCTGTACCTCTTGGTCTTTGAATAAAGCCTGAGTAGGAAAAAAAAAAAAAAAAAAAAAAAAAAAAAAAAAAAAAAAAAAAAAAAAAAAAAAAAAAAAAAAAAAAAAAAAAAAAAAAAAAAAAAAAAAAAAAAAAAAAAAAAAAAAAAAAAAAAAAAA]

- **1xMS2(U)site2-Luc2**

T7 promoter: 11-30

MS2-binding motif (U-variant): 55-73

Kozak sequence (including start codon): 98-106

Luc2: 107-1753

Stop codon: 1754-1756

[CAGTGAATTGTAATACGACTCACTATAGGGCGAATTAAGAGAGAAAAGAAGAGTACATGAGGATTACCCATGTAAGAAGAAATATAAGACACCGGTCGCCACCATGGAAGATGCCAAAAACATTAAGAAGGGCCCAGCGCCATTCTACCCACTCGAAGACGGGACCGCCGGCGAGCAGCTGCACAAAGCCATGAAGCGCTACGCCCTGGTGCCCGGCACCATCGCCTTTACCGACGCACATATCGAGGTGGACATTACCTACGCCGAGTACTTCGAGATGAGCGTTCGGCTGGCAGAAGCTATGAAGCGCTATGGGCTGAATACAAACCATCGGATCGTGGTGTGCAGCGAGAATAGCTTGCAGTTCTTCATGCCCGTGTTGGGTGCCCTGTTCATCGGTGTGGCTGTGGCCCCAGCTAACGACATCTACAACGAGCGCGAGCTGCTGAACAGCATGGGCATCAGCCAGCCCACCGTCGTATTCGTGAGCAAGAAAGGGCTGCAAAAGATCCTCAACGTGCAAAAGAAGCTACCGATCATACAAAAGATCATCATCATGGATAGCAAGACCGACTACCAGGGCTTCCAAAGCATGTACACCTTCGTGACTTCCCATTTGCCACCCGGCTTCAACGAGTACGACTTCGTGCCCGAGAGCTTCGACCGGGACAAAACCATCGCCCTGATCATGAACAGTAGTGGCAGTACCGGATTGCCCAAGGGCGTAGCCCTACCGCACCGCACCGCTTGTGTCCGATTCAGTCATGCCCGCGACCCCATCTTCGGCAACCAGATCATCCCCGACACCGCTATCCTCAGCGTGGTGCCATTTCACCACGGCTTCGGCATGTTCACCACGCTGGGCTACTTGATCTGCGGCTTTCGGGTCGTGCTCATGTACCGCTTCGAGGAGGAGCTATTCTTGCGCAGCTTGCAAGACTATAAGATTCAATCTGCCCTGCTGGTGCCCACACTATTTAGCTTCTTCGCTAAGAGCACTCTCATCGACAAGTACGACCTAAGCAACTTGCACGAGATCGCCAGCGGCGGGGCGCCGCTCAGCAAGGAGGTAGGTGAGGCCGTGGCCAAACGCTTCCACCTACCAGGCATCCGCCAGGGCTACGGCCTGACAGAAACAACCAGCGCCATTCTGATCACCCCCGAAGGGGACGACAAGCCTGGCGCAGTAGGCAAGGTGGTGCCCTTCTTCGAGGCTAAGGTGGTGGACTTGGACACCGGTAAGACACTGGGTGTGAACCAGCGCGGCGAGCTGTGCGTCCGTGGCCCCATGATCATGAGCGGCTACGTTAACAACCCCGAGGCTACAAACGCTCTCATCGACAAGGACGGCTGGCTGCACAGCGGCGACATCGCCTACTGGGACGAGGACGAGCACTTCTTCATCGTGGACCGGCTGAAGAGCCTGATCAAATACAAGGGCTACCAGGTAGCCCCAGCCGAACTGGAGAGCATCCTGCTGCAACACCCCAACATCTTCGACGCCGGGGTCGCCGGCCTGCCCGACGACGATGCCGGCGAGCTGCCCGCCGCAGTCGTCGTGCTGGAACACGGTAAAACCATGACCGAGAAGGAGATCGTGGACTATGTGGCCAGCCAGGTTACAACCGCCAAGAAGCTGCGCGGTGGTGTTGTGTTCGTGGACGAGGTGCCTAAAGGACTGACCGGCAAGTTGGACGCCCGCAAGATCCGCGAGATTCTCATTAAGGCCAAGAAGGGCGGCAAGATCGCCGTGTAAATCTAGACCTTCTGCGGGGCTTGCCTTCTGGCCATGCCCTTCTTCTCTCCCTTGCACCTGTACCTCTTGGTCTTTGAATAAAGCCTGAGTAGGAAAAAAAAAAAAAAAAAAAAAAAAAAAAAAAAAAAAAAAAAAAAAAAAAAAAAAAAAAAAAAAAAAAAAAAAAAAAAAAAAAAAAAAAAAAAAAAAAAAAAAAAAAAAAAAAAAAAAAAA]

- **Nluc**

T7 promoter (for CleanCap AG Reagent): 11-30

Kozak sequence (including start codon): 79-87

Nluc: 88-597

Stop codon: 598-600

[CAGTGAATTGTAATACGACTCACTATAAGGCGAATTAAGAGAGAAAAGAAGAGTAAGAAGAAATATAAGACACCGGTCGCCACCATGGTCTTCACACTCGAAGATTTCGTTGGGGACTGGCGACAGACAGCCGGCTACAACCTGGACCAAGTCCTTGAACAGGGAGGTGTGTCCAGTTTGTTTCAGAATCTCGGGGTGTCCGTAACTCCGATCCAAAGGATTGTCCTGAGCGGTGAAAATGGGCTGAAGATCGACATCCATGTCATCATCCCGTATGAAGGTCTGAGCGGCGACCAAATGGGCCAGATCGAAAAAATTTTTAAGGTGGTGTACCCTGTGGATGATCATCACTTTAAGGTGATCCTGCACTATGGCACACTGGTAATCGACGGGGTTACGCCGAACATGATCGACTATTTCGGACGGCCGTATGAAGGCATCGCCGTGTTCGACGGCAAAAAGATCACTGTAACAGGGACCCTGTGGAACGGCAACAAAATTATCGACGAGCGCCTGATCAACCCCGACGGCTCCCTGCTGTTCCGAGTAACCATCAACGGAGTGACCGGCTGGCGGCTGTGCGAACGCATTCTGGCGTAAATCTAGACCTTCTGCGGGGCTTGCCTTCTGGCCATGCCCTTCTTCTCTCCCTTGCACCTGTACCTCTTGGTCTTTGAATAAAGCCTGAGTAGGAAAAAAAAAAAAAAAAAAAAAAAAAAAAAAAAAAAAAAAAAAAAAAAAAAAAAAAAAAAAAAAAAAAAAAAAAAAAAAAAAAAAAAAAAAAAAAAAAAAAAAAAAAAAAAAAAAAAAAAA]

- **ARRB2-tevD-CaVT**

T7 promoter (for CleanCap AG Reagent): 11-30

Kozak sequence (including start codon): 79-87

ARRB2: 88-1311

3xGGGGS flexible linker: 1312-1356

tevD: 1357-1419

MS2CP: 1420-1767

FCV-derived VPg: 1774-2106

Stop codon: 2107-2109

[CAGTGAATTGTAATACGACTCACTATAAGGCGAATTAAGAGAGAAAAGAAGAGTAAGAAGAAATATAAGACACCGGTCGCCACCATGGGAGAAAAGCCTGGCACCAGAGTGTTCAAGAAGTCTAGCCCCAACTGCAAGCTGACCGTGTACCTCGGCAAGCGGGACTTCGTGGACCACCTGGACAAAGTGGATCCCGTGGATGGCGTGGTGCTGGTGGATCCTGACTACCTGAAGGACCGGAAGGTGTTCGTGACCCTGACCTGCGCCTTCAGATACGGCAGAGAAGATCTGGACGTGCTGGGCCTGAGCTTCAGAAAGGACCTGTTTATCGCCACCTACCAGGCCTTTCCACCTGTGCCTAATCCTCCACGGCCTCCTACCAGACTGCAGGACAGACTGCTGAGAAAGCTGGGCCAGCACGCTCACCCCTTTTTCTTCACAATCCCTCAAAACCTGCCTTGCAGCGTGACACTGCAGCCCGGACCTGAGGATACAGGCAAAGCTTGCGGCGTGGACTTCGAGATCAGAGCCTTCTGTGCCAAGAGCCTGGAAGAGAAGTCCCACAAGAGAAACAGCGTGCGGCTGGTCATCAGAAAGGTGCAGTTCGCCCCTGAGAAGCCTGGACCTCAGCCTTCTGCCGAGACAACCAGACACTTCCTGATGAGCGACCGCAGCCTGCATCTGGAAGCCAGCCTCGACAAAGAGCTGTACTACCACGGCGAGCCCCTGAACGTCAACGTCCACGTGACCAACAACAGCACCAAGACCGTGAAAAAGATCAAGGTGTCCGTGCGGCAGTACGCCGACATCTGCCTGTTTAGCACAGCCCAGTACAAGTGCCCCGTGGCTCAGCTGGAACAGGACGATCAAGTGTCCCCTAGCAGCACCTTCTGCAAGGTGTACACAATCACCCCTCTGCTGAGCGACAACAGAGAGAAGAGAGGACTGGCCCTGGACGGCAAGCTGAAGCACGAGGATACAAACCTGGCCAGCAGCACCATCGTGAAAGAGGGCGCCAACAAAGAGGTGCTGGGCATCCTGGTGTCCTACAGAGTGAAAGTGAAGCTGGTGGTGTCCAGAGGCGGCGACGTTTCAGTGGAACTGCCCTTCGTGCTGATGCACCCCAAGCCTCACGATCACATCCCTCTGCCTAGACCTCAGAGCGCCGCTCCTGAAACAGATGTGCCCGTGGACACCAACCTGATCGAGTTCGACACCAACTACGCCACCGACGACGACATCGTGTTCGAGGACTTCGCCCGGCTGAGACTGAAGGGCATGAAGGACGACGACTACGACGATCAGCTTTGTGGCGGCGGAGGATCTGGCGGAGGTGGAAGCGGAGGCGGTGGATCTGAGAACCTGTACTTCCAGTTCCACAAGAGCGGCGCCTGGAAGCTGCCTGTGTCTCTGGTTAAGGCTTCTAACTTTACTCAGTTCGTTCTCGTCGACAATGGCGGAACTGGCGACGTGACTGTCGCCCCAAGCAACTTCGCTAACGGGGTCGCTGAATGGATCAGCTCTAACTCGCGATCACAGGCTTACAAAGTAACCTGTAGCGTTCGTCAGAGCTCTGCGCAGAATCGCAAATACACCATCAAAGTCGAGGTGCCTAAAGGCGCATGGAGGTCTTACTTAAATATGGAACTAACCATTCCAATTTTCGCCACGAATTCCGACTGCGAGCTTATTGTTAAGGCAATGCAAGGTCTCCTAAAAGATGGAAACCCGATTCCCTCGGCCATCGCGGCCAACTCCGGCATCTACGGATCCGCCAAGGGCAAGACCAAGAGCAAAGTGGGCCCCTACAGAGGCAGAGGCGTGGCCCTTACAGACGACGAGTATGACGAATGGCGCGAGCACAACGCCACCAGAAAGCTGGATCTGAGCGTGGAAGATTTCCTGATGCTGCGGCACAGAGCCGCTCTGGGAGCTGATGATGCCGACGCCGTGAAGTTCAGATCCTGGTGGAACAGCAGAAGCCGGCTGGCCGACGATTACGAGGATGTGACCGTGATCGGCAAAGGCGGCGTGAAGCACGAGAAGATCCGGACCAATACTCTGAGAGCCGTGGACAGAGGCTACGACGTGTCCTTCGCTGAAGAATGAATCTAGACCTTCTGCGGGGCTTGCCTTCTGGCCATGCCCTTCTTCTCTCCCTTGCACCTGTACCTCTTGGTCTTTGAATAAAGCCTGAGTAGGAAAAAAAAAAAAAAAAAAAAAAAAAAAAAAAAAAAAAAAAAAAAAAAAAAAAAAAAAAAAAAAAAAAAAAAAAAAAAAAAAAAAAAAAAAAAAAAAAAAAAAAAAAAAAAAAAAAAAAAA]

- **ARRB2-tevD-CaVT_ver.2**

T7 promoter (for CleanCap AG Reagent): 11-30

Kozak sequence (including start codon): 79-87

ARRB2: 88-1311

3xGGGGS flexible linker: 1312-1356

tevD: 1357-1419

2xGGGGS flexible linker: 1420-1449

MS2CP: 1450-1797

FCV-derived VPg: 1804-2136

Stop codon: 2137-2139

[CAGTGAATTGTAATACGACTCACTATAAGGCGAATTAAGAGAGAAAAGAAGAGTAAGAAGAAATATAAGACACCGGTCGCCACCATGGGAGAAAAGCCTGGCACCAGAGTGTTCAAGAAGTCTAGCCCCAACTGCAAGCTGACCGTGTACCTCGGCAAGCGGGACTTCGTGGACCACCTGGACAAAGTGGATCCCGTGGATGGCGTGGTGCTGGTGGATCCTGACTACCTGAAGGACCGGAAGGTGTTCGTGACCCTGACCTGCGCCTTCAGATACGGCAGAGAAGATCTGGACGTGCTGGGCCTGAGCTTCAGAAAGGACCTGTTTATCGCCACCTACCAGGCCTTTCCACCTGTGCCTAATCCTCCACGGCCTCCTACCAGACTGCAGGACAGACTGCTGAGAAAGCTGGGCCAGCACGCTCACCCCTTTTTCTTCACAATCCCTCAAAACCTGCCTTGCAGCGTGACACTGCAGCCCGGACCTGAGGATACAGGCAAAGCTTGCGGCGTGGACTTCGAGATCAGAGCCTTCTGTGCCAAGAGCCTGGAAGAGAAGTCCCACAAGAGAAACAGCGTGCGGCTGGTCATCAGAAAGGTGCAGTTCGCCCCTGAGAAGCCTGGACCTCAGCCTTCTGCCGAGACAACCAGACACTTCCTGATGAGCGACCGCAGCCTGCATCTGGAAGCCAGCCTCGACAAAGAGCTGTACTACCACGGCGAGCCCCTGAACGTCAACGTCCACGTGACCAACAACAGCACCAAGACCGTGAAAAAGATCAAGGTGTCCGTGCGGCAGTACGCCGACATCTGCCTGTTTAGCACAGCCCAGTACAAGTGCCCCGTGGCTCAGCTGGAACAGGACGATCAAGTGTCCCCTAGCAGCACCTTCTGCAAGGTGTACACAATCACCCCTCTGCTGAGCGACAACAGAGAGAAGAGAGGACTGGCCCTGGACGGCAAGCTGAAGCACGAGGATACAAACCTGGCCAGCAGCACCATCGTGAAAGAGGGCGCCAACAAAGAGGTGCTGGGCATCCTGGTGTCCTACAGAGTGAAAGTGAAGCTGGTGGTGTCCAGAGGCGGCGACGTTTCAGTGGAACTGCCCTTCGTGCTGATGCACCCCAAGCCTCACGATCACATCCCTCTGCCTAGACCTCAGAGCGCCGCTCCTGAAACAGATGTGCCCGTGGACACCAACCTGATCGAGTTCGACACCAACTACGCCACCGACGACGACATCGTGTTCGAGGACTTCGCCCGGCTGAGACTGAAGGGCATGAAGGACGACGACTACGACGATCAGCTTTGTGGCGGCGGAGGATCTGGCGGAGGTGGAAGCGGAGGCGGTGGATCTGAGAACCTGTACTTCCAGTTCCACAAGAGCGGCGCCTGGAAGCTGCCTGTGTCTCTGGTTAAGGGCGGCGGCGGCTCCGGCGGCGGCGGCTCCGCTTCTAACTTTACTCAGTTCGTTCTCGTCGACAATGGCGGAACTGGCGACGTGACTGTCGCCCCAAGCAACTTCGCTAACGGGGTCGCTGAATGGATCAGCTCTAACTCGCGATCACAGGCTTACAAAGTAACCTGTAGCGTTCGTCAGAGCTCTGCGCAGAATCGCAAATACACCATCAAAGTCGAGGTGCCTAAAGGCGCATGGAGGTCTTACTTAAATATGGAACTAACCATTCCAATTTTCGCCACGAATTCCGACTGCGAGCTTATTGTTAAGGCAATGCAAGGTCTCCTAAAAGATGGAAACCCGATTCCCTCGGCCATCGCGGCCAACTCCGGCATCTACGGATCCGCCAAGGGCAAGACCAAGAGCAAAGTGGGCCCCTACAGAGGCAGAGGCGTGGCCCTTACAGACGACGAGTATGACGAATGGCGCGAGCACAACGCCACCAGAAAGCTGGATCTGAGCGTGGAAGATTTCCTGATGCTGCGGCACAGAGCCGCTCTGGGAGCTGATGATGCCGACGCCGTGAAGTTCAGATCCTGGTGGAACAGCAGAAGCCGGCTGGCCGACGATTACGAGGATGTGACCGTGATCGGCAAAGGCGGCGTGAAGCACGAGAAGATCCGGACCAATACTCTGAGAGCCGTGGACAGAGGCTACGACGTGTCCTTCGCTGAAGAATGAATCTAGACCTTCTGCGGGGCTTGCCTTCTGGCCATGCCCTTCTTCTCTCCCTTGCACCTGTACCTCTTGGTCTTTGAATAAAGCCTGAGTAGGAAAAAAAAAAAAAAAAAAAAAAAAAAAAAAAAAAAAAAAAAAAAAAAAAAAAAAAAAAAAAAAAAAAAAAAAAAAAAAAAAAAAAAAAAAAAAAAAAAAAAAAAAAAAAAAAAAAAAAAA]

- **EGFP**

T7 promoter (for CleanCap AG Reagent): 11-30

Kozak sequence (including start codon): 79-87

EGFP: 88-801

Stop codon: 802-804

[CAGTGAATTGTAATACGACTCACTATAAGGCGAATTAAGAGAGAAAAGAAGAGTAAGAAGAAATATAAGACACCGGTCGCCACCATGGTGAGCAAGGGCGAGGAGCTGTTCACCGGGGTGGTGCCCATCCTGGTCGAGCTGGACGGCGACGTAAACGGCCACAAGTTCAGCGTGTCCGGCGAGGGCGAGGGCGATGCCACCTACGGCAAGCTGACCCTGAAGTTCATCTGCACCACCGGCAAGCTGCCCGTGCCCTGGCCCACCCTCGTGACCACCCTGACCTACGGCGTGCAGTGCTTCAGCCGCTACCCCGACCACATGAAGCAGCACGACTTCTTCAAGTCCGCCATGCCCGAAGGCTACGTCCAGGAGCGCACCATCTTCTTCAAGGACGACGGCAACTACAAGACCCGCGCCGAGGTGAAGTTCGAGGGCGACACCCTGGTGAACCGCATCGAGCTGAAGGGCATCGACTTCAAGGAGGACGGCAACATCCTGGGGCACAAGCTGGAGTACAACTACAACAGCCACAACGTCTATATCATGGCCGACAAGCAGAAGAACGGCATCAAGGTGAACTTCAAGATCCGCCACAACATCGAGGACGGCAGCGTGCAGCTCGCCGACCACTACCAGCAGAACACCCCCATCGGCGACGGCCCCGTGCTGCTGCCCGACAACCACTACCTGAGCACCCAGTCCGCCCTGAGCAAAGACCCCAACGAGAAGCGCGATCACATGGTCCTGCTGGAGTTCGTGACCGCCGCCGGGATCACTCTCGGCATGGACGAGCTGTACAAGTAAATCTAGACCTTCTGCGGGGCTTGCCTTCTGGCCATGCCCTTCTTCTCTCCCTTGCACCTGTACCTCTTGGTCTTTGAATAAAGCCTGAGTAGGAAAAAAAAAAAAAAAAAAAAAAAAAAAAAAAAAAAAAAAAAAAAAAAAAAAAAAAAAAAAAAAAAAAAAAAAAAAAAAAAAAAAAAAAAAAAAAAAAAAAAAAAAAAAAAAAAAAAAAAA]

- **CaVT**

T7 promoter (for CleanCap AG Reagent): 11-30

Kozak sequence (including start codon): 79-87

MS2CP: 88-435

FCV-derived VPg: 442-774

Stop codon: 775-777

[CAGTGAATTGTAATACGACTCACTATAAGGCGAATTAAGAGAGAAAAGAAGAGTAAGAAGAAATATAAGACACCGGTCGCCACCATGGCTTCTAACTTTACTCAGTTCGTTCTCGTCGACAATGGCGGAACTGGCGACGTGACTGTCGCCCCAAGCAACTTCGCTAACGGGGTCGCTGAATGGATCAGCTCTAACTCGCGATCACAGGCTTACAAAGTAACCTGTAGCGTTCGTCAGAGCTCTGCGCAGAATCGCAAATACACCATCAAAGTCGAGGTGCCTAAAGGCGCATGGAGGTCTTACTTAAATATGGAACTAACCATTCCAATTTTCGCCACGAATTCCGACTGCGAGCTTATTGTTAAGGCAATGCAAGGTCTCCTAAAAGATGGAAACCCGATTCCCTCGGCCATCGCGGCCAACTCCGGCATCTACGGATCCGCCAAGGGCAAGACCAAGAGCAAAGTGGGCCCCTACAGAGGCAGAGGCGTGGCCCTTACAGACGACGAGTATGACGAATGGCGCGAGCACAACGCCACCAGAAAGCTGGATCTGAGCGTGGAAGATTTCCTGATGCTGCGGCACAGAGCCGCTCTGGGAGCTGATGATGCCGACGCCGTGAAGTTCAGATCCTGGTGGAACAGCAGAAGCCGGCTGGCCGACGATTACGAGGATGTGACCGTGATCGGCAAAGGCGGCGTGAAGCACGAGAAGATCCGGACCAATACTCTGAGAGCCGTGGACAGAGGCTACGACGTGTCCTTCGCTGAAGAATGAATCTAGACCTTCTGCGGGGCTTGCCTTCTGGCCATGCCCTTCTTCTCTCCCTTGCACCTGTACCTCTTGGTCTTTGAATAAAGCCTGAGTAGGAAAAAAAAAAAAAAAAAAAAAAAAAAAAAAAAAAAAAAAAAAAAAAAAAAAAAAAAAAAAAAAAAAAAAAAAAAAAAAAAAAAAAAAAAAAAAAAAAAAAAAAAAAAAAAAAAAAAAAAA]

- **TEVp**

T7 promoter (for CleanCap AG Reagent): 11-30

Kozak sequence (including start codon): 79-87

TEV protease (S219V mutant): 88-792

Stop codon: 793-795

[CAGTGAATTGTAATACGACTCACTATAAGGCGAATTAAGAGAGAAAAGAAGAGTAAGAAGAAATATAAGACACCGGTCGCCACCATGGAGAGCCTGTTTAAGGGCCCCAGAGACTACAACCCCATCAGCTCCACCATCTGCCACCTGACCAATGAGTCCGATGGCCACACCACAAGCCTGTACGGCATCGGCTTCGGCCCCTTCATCATCACCAACAAGCACCTGTTCAGACGGAACAACGGCACCCTGCTGGTGCAGTCTCTGCACGGCGTGTTCAAAGTGAAGAATACCACCACACTGCAGCAGCACCTGATCGACGGCCGGGACATGATCATCATCAGAATGCCCAAGGACTTCCCGCCTTTTCCACAGAAGCTGAAGTTCAGAGAGCCCCAGCGCGAGGAAAGAATCTGCCTGGTCACCACCAACTTCCAGACCAAGAGCATGTCCTCCATGGTGTCCGATACCAGCTGCACATTCCCTAGCAGCGACGGCATCTTCTGGAAGCACTGGATTCAGACCAAGGACGGCCAGTGTGGCAGCCCTCTGGTGTCTACAAGAGATGGCTTCATCGTGGGCATCCACAGCGCCAGCAACTTCACCAATACCAACAACTACTTCACCAGCGTGCCGAAGAACTTCATGGAACTGCTGACCAATCAAGAGGCTCAGCAGTGGGTTTCCGGCTGGCGGCTGAATGCTGATTCTGTGCTGTGGGGCGGACACAAGGTGTTCATGGTCAAGCCCGAGGAACCCTTCCAGCCTGTGAAAGAGGCCACACAGCTGATGAACTGAATCTAGACCTTCTGCGGGGCTTGCCTTCTGGCCATGCCCTTCTTCTCTCCCTTGCACCTGTACCTCTTGGTCTTTGAATAAAGCCTGAGTAGGAAAAAAAAAAAAAAAAAAAAAAAAAAAAAAAAAAAAAAAAAAAAAAAAAAAAAAAAAAAAAAAAAAAAAAAAAAAAAAAAAAAAAAAAAAAAAAAAAAAAAAAAAAAAAAAAAAAAAAAA]

- **TEVp (C151A)**

T7 promoter (for CleanCap AG Reagent): 11-30

Kozak sequence (including start codon): 79-87

TEV protease (C151A/S219V mutant): 88-792

Stop codon: 793-795

[CAGTGAATTGTAATACGACTCACTATAAGGCGAATTAAGAGAGAAAAGAAGAGTAAGAAGAAATATAAGACACCGGTCGCCACCATGGAGAGCCTGTTTAAGGGCCCCAGAGACTACAACCCCATCAGCTCCACCATCTGCCACCTGACCAATGAGTCCGATGGCCACACCACAAGCCTGTACGGCATCGGCTTCGGCCCCTTCATCATCACCAACAAGCACCTGTTCAGACGGAACAACGGCACCCTGCTGGTGCAGTCTCTGCACGGCGTGTTCAAAGTGAAGAATACCACCACACTGCAGCAGCACCTGATCGACGGCCGGGACATGATCATCATCAGAATGCCCAAGGACTTCCCGCCTTTTCCACAGAAGCTGAAGTTCAGAGAGCCCCAGCGCGAGGAAAGAATCTGCCTGGTCACCACCAACTTCCAGACCAAGAGCATGTCCTCCATGGTGTCCGATACCAGCTGCACATTCCCTAGCAGCGACGGCATCTTCTGGAAGCACTGGATTCAGACCAAGGACGGCCAGGCCGGCAGCCCTCTGGTGTCTACAAGAGATGGCTTCATCGTGGGCATCCACAGCGCCAGCAACTTCACCAATACCAACAACTACTTCACCAGCGTGCCGAAGAACTTCATGGAACTGCTGACCAATCAAGAGGCTCAGCAGTGGGTTTCCGGCTGGCGGCTGAATGCTGATTCTGTGCTGTGGGGCGGACACAAGGTGTTCATGGTCAAGCCCGAGGAACCCTTCCAGCCTGTGAAAGAGGCCACACAGCTGATGAACTGAATCTAGACCTTCTGCGGGGCTTGCCTTCTGGCCATGCCCTTCTTCTCTCCCTTGCACCTGTACCTCTTGGTCTTTGAATAAAGCCTGAGTAGGAAAAAAAAAAAAAAAAAAAAAAAAAAAAAAAAAAAAAAAAAAAAAAAAAAAAAAAAAAAAAAAAAAAAAAAAAAAAAAAAAAAAAAAAAAAAAAAAAAAAAAAAAAAAAAAAAAAAAAAA]

- **AVPR2-TEVp**

T7 promoter (for CleanCap AG Reagent): 11-30

Kozak sequence (including start codon): 79-87

Human AVPR2: 88-1197

3xGGGGS flexible linker: 1198-1242

TEV protease (S219V mutant): 1243-1947

Stop codon: 1948-1950

[CAGTGAATTGTAATACGACTCACTATAAGGCGAATTAAGAGAGAAAAGAAGAGTAAGAAGAAATATAAGACACCGGTCGCCACCATGCTGATGGCCTCTACAACATCTGCCGTGCCTGGACACCCTAGCCTGCCTTCTCTGCCTAGCAACAGCAGCCAAGAGAGGCCCCTGGATACCAGAGATCCTCTGCTGGCCAGAGCCGAACTGGCCCTGCTGTCTATCGTGTTTGTGGCCGTGGCTCTGTCCAACGGACTGGTTCTTGCTGCCCTGGCTCGGAGAGGAAGAAGAGGACATTGGGCCCCTATCCACGTGTTCATCGGCCATCTGTGTCTGGCCGATCTGGCTGTGGCACTGTTTCAGGTTCTGCCTCAGCTGGCCTGGAAGGCCACCGATAGATTCAGAGGCCCCGACGCTCTGTGCAGAGCCGTGAAATACCTGCAGATGGTCGGGATGTACGCCAGCAGCTACATGATCCTGGCCATGACACTGGACCGGCACAGAGCCATCTGTAGACCCATGCTGGCCTACAGACATGGCTCTGGCGCCCACTGGAATAGACCTGTGCTTGTGGCCTGGGCCTTCAGCCTGCTTCTGTCTCTGCCCCAGCTGTTCATCTTCGCCCAGAGAAATGTGGAAGGCGGCAGCGGCGTTACAGATTGCTGGGCCTGTTTTGCCGAGCCTTGGGGTAGAAGAACCTACGTGACATGGATCGCCCTGATGGTGTTCGTGGCCCCTACACTGGGAATTGCCGCTTGCCAGGTGCTGATCTTCAGAGAGATCCACGCCAGCCTGGTGCCAGGACCTTCTGAAAGACCTGGCGGACGCAGACGGGGAAGAAGAACAGGATCTCCTGGCGAAGGCGCCCATGTGTCTGCTGCCGTGGCCAAGACAGTGCGGATGACACTCGTGATCGTGGTGGTGTACGTGCTGTGCTGGGCCCCATTCTTTCTGGTGCAACTGTGGGCCGCCTGGGATCCTGAAGCTCCTCTTGAAGGCGCTCCCTTCGTGCTGCTGATGCTGCTGGCTAGCCTGAACTCCTGCACAAACCCCTGGATCTACGCCTCCTTCAGCAGCAGCGTGTCCAGCGAGCTGAGAAGCCTGCTGTGTTGTGCCAGAGGCAGGACACCTCCATCTCTGGGACCTCAGGATGAGAGCTGTACCACCGCCTCTTCTAGCCTGGCCAAGGATACAAGTTCTGGCGGCGGAGGATCTGGCGGAGGTGGAAGCGGAGGCGGCGGATCTGAGAGCCTGTTTAAGGGCCCCAGAGACTACAACCCCATCAGCTCCACCATCTGCCACCTGACCAATGAGTCCGATGGCCACACCACAAGCCTGTACGGCATCGGCTTCGGCCCCTTCATCATCACCAACAAGCACCTGTTCAGACGGAACAACGGCACCCTGCTGGTGCAGTCTCTGCACGGCGTGTTCAAAGTGAAGAATACCACCACACTGCAGCAGCACCTGATCGACGGCCGGGACATGATCATCATCAGAATGCCCAAGGACTTCCCGCCTTTTCCACAGAAGCTGAAGTTCAGAGAGCCCCAGCGCGAGGAAAGAATCTGCCTGGTCACCACCAACTTCCAGACCAAGAGCATGTCCTCCATGGTGTCCGATACCAGCTGCACATTCCCTAGCAGCGACGGCATCTTCTGGAAGCACTGGATTCAGACCAAGGACGGCCAGTGTGGCAGCCCTCTGGTGTCTACAAGAGATGGCTTCATCGTGGGCATCCACAGCGCCAGCAACTTCACCAATACCAACAACTACTTCACCAGCGTGCCGAAGAACTTCATGGAACTGCTGACCAATCAAGAGGCTCAGCAGTGGGTTTCCGGCTGGCGGCTGAATGCTGATTCTGTGCTGTGGGGCGGACACAAGGTGTTCATGGTCAAGCCCGAGGAACCCTTCCAGCCTGTGAAAGAGGCCACACAGCTGATGAACTGAATCTAGACCTTCTGCGGGGCTTGCCTTCTGGCCATGCCCTTCTTCTCTCCCTTGCACCTGTACCTCTTGGTCTTTGAATAAAGCCTGAGTAGGAAAAAAAAAAAAAAAAAAAAAAAAAAAAAAAAAAAAAAAAAAAAAAAAAAAAAAAAAAAAAAAAAAAAAAAAAAAAAAAAAAAAAAAAAAAAAAAAAAAAAAAAAAAAAAAAAAAAAAAA]

- **AVPR2-TEVpΔ**

T7 promoter (for CleanCap AG Reagent): 11-30

Kozak sequence (including start codon): 79-87

Human AVPR2: 88-1197

3xGGGGS flexible linker: 1198-1242

Carboxy-terminally truncated TEVp (S219V mutant): 1243-1896

Stop codon: 1897-1899

[CAGTGAATTGTAATACGACTCACTATAAGGCGAATTAAGAGAGAAAAGAAGAGTAAGAAGAAATATAAGACACCGGTCGCCACCATGCTGATGGCCTCTACAACATCTGCCGTGCCTGGACACCCTAGCCTGCCTTCTCTGCCTAGCAACAGCAGCCAAGAGAGGCCCCTGGATACCAGAGATCCTCTGCTGGCCAGAGCCGAACTGGCCCTGCTGTCTATCGTGTTTGTGGCCGTGGCTCTGTCCAACGGACTGGTTCTTGCTGCCCTGGCTCGGAGAGGAAGAAGAGGACATTGGGCCCCTATCCACGTGTTCATCGGCCATCTGTGTCTGGCCGATCTGGCTGTGGCACTGTTTCAGGTTCTGCCTCAGCTGGCCTGGAAGGCCACCGATAGATTCAGAGGCCCCGACGCTCTGTGCAGAGCCGTGAAATACCTGCAGATGGTCGGGATGTACGCCAGCAGCTACATGATCCTGGCCATGACACTGGACCGGCACAGAGCCATCTGTAGACCCATGCTGGCCTACAGACATGGCTCTGGCGCCCACTGGAATAGACCTGTGCTTGTGGCCTGGGCCTTCAGCCTGCTTCTGTCTCTGCCCCAGCTGTTCATCTTCGCCCAGAGAAATGTGGAAGGCGGCAGCGGCGTTACAGATTGCTGGGCCTGTTTTGCCGAGCCTTGGGGTAGAAGAACCTACGTGACATGGATCGCCCTGATGGTGTTCGTGGCCCCTACACTGGGAATTGCCGCTTGCCAGGTGCTGATCTTCAGAGAGATCCACGCCAGCCTGGTGCCAGGACCTTCTGAAAGACCTGGCGGACGCAGACGGGGAAGAAGAACAGGATCTCCTGGCGAAGGCGCCCATGTGTCTGCTGCCGTGGCCAAGACAGTGCGGATGACACTCGTGATCGTGGTGGTGTACGTGCTGTGCTGGGCCCCATTCTTTCTGGTGCAACTGTGGGCCGCCTGGGATCCTGAAGCTCCTCTTGAAGGCGCTCCCTTCGTGCTGCTGATGCTGCTGGCTAGCCTGAACTCCTGCACAAACCCCTGGATCTACGCCTCCTTCAGCAGCAGCGTGTCCAGCGAGCTGAGAAGCCTGCTGTGTTGTGCCAGAGGCAGGACACCTCCATCTCTGGGACCTCAGGATGAGAGCTGTACCACCGCCTCTTCTAGCCTGGCCAAGGATACAAGTTCTGGCGGCGGAGGATCTGGCGGAGGTGGAAGCGGAGGCGGCGGATCTGAGAGCCTGTTTAAGGGCCCCAGAGACTACAACCCCATCAGCTCCACCATCTGCCACCTGACCAATGAGTCCGATGGCCACACCACAAGCCTGTACGGCATCGGCTTCGGCCCCTTCATCATCACCAACAAGCACCTGTTCAGACGGAACAACGGCACCCTGCTGGTGCAGTCTCTGCACGGCGTGTTCAAAGTGAAGAATACCACCACACTGCAGCAGCACCTGATCGACGGCCGGGACATGATCATCATCAGAATGCCCAAGGACTTCCCGCCTTTTCCACAGAAGCTGAAGTTCAGAGAGCCCCAGCGCGAGGAAAGAATCTGCCTGGTCACCACCAACTTCCAGACCAAGAGCATGTCCTCCATGGTGTCCGATACCAGCTGCACATTCCCTAGCAGCGACGGCATCTTCTGGAAGCACTGGATTCAGACCAAGGACGGCCAGTGTGGCAGCCCTCTGGTGTCTACAAGAGATGGCTTCATCGTGGGCATCCACAGCGCCAGCAACTTCACCAATACCAACAACTACTTCACCAGCGTGCCGAAGAACTTCATGGAACTGCTGACCAATCAAGAGGCTCAGCAGTGGGTTTCCGGCTGGCGGCTGAATGCTGATTCTGTGCTGTGGGGCGGACACAAGGTGTTCATGGTCTGAATCTAGACCTTCTGCGGGGCTTGCCTTCTGGCCATGCCCTTCTTCTCTCCCTTGCACCTGTACCTCTTGGTCTTTGAATAAAGCCTGAGTAGGAAAAAAAAAAAAAAAAAAAAAAAAAAAAAAAAAAAAAAAAAAAAAAAAAAAAAAAAAAAAAAAAAAAAAAAAAAAAAAAAAAAAAAAAAAAAAAAAAAAAAAAAAAAAAAAAAAAAAAAA]

- **AVPR2-TEVpΔT30A**

T7 promoter (for CleanCap AG Reagent): 11-30

Kozak sequence (including start codon): 79-87

Human AVPR2: 88-1197

3xGGGGS flexible linker: 1198-1242

Carboxy-terminally truncated TEVp (T30A/S219V mutant): 1243-1896

Stop codon: 1897-1899

[CAGTGAATTGTAATACGACTCACTATAAGGCGAATTAAGAGAGAAAAGAAGAGTAAGAAGAAATATAAGACACCGGTCGCCACCATGCTGATGGCCTCTACAACATCTGCCGTGCCTGGACACCCTAGCCTGCCTTCTCTGCCTAGCAACAGCAGCCAAGAGAGGCCCCTGGATACCAGAGATCCTCTGCTGGCCAGAGCCGAACTGGCCCTGCTGTCTATCGTGTTTGTGGCCGTGGCTCTGTCCAACGGACTGGTTCTTGCTGCCCTGGCTCGGAGAGGAAGAAGAGGACATTGGGCCCCTATCCACGTGTTCATCGGCCATCTGTGTCTGGCCGATCTGGCTGTGGCACTGTTTCAGGTTCTGCCTCAGCTGGCCTGGAAGGCCACCGATAGATTCAGAGGCCCCGACGCTCTGTGCAGAGCCGTGAAATACCTGCAGATGGTCGGGATGTACGCCAGCAGCTACATGATCCTGGCCATGACACTGGACCGGCACAGAGCCATCTGTAGACCCATGCTGGCCTACAGACATGGCTCTGGCGCCCACTGGAATAGACCTGTGCTTGTGGCCTGGGCCTTCAGCCTGCTTCTGTCTCTGCCCCAGCTGTTCATCTTCGCCCAGAGAAATGTGGAAGGCGGCAGCGGCGTTACAGATTGCTGGGCCTGTTTTGCCGAGCCTTGGGGTAGAAGAACCTACGTGACATGGATCGCCCTGATGGTGTTCGTGGCCCCTACACTGGGAATTGCCGCTTGCCAGGTGCTGATCTTCAGAGAGATCCACGCCAGCCTGGTGCCAGGACCTTCTGAAAGACCTGGCGGACGCAGACGGGGAAGAAGAACAGGATCTCCTGGCGAAGGCGCCCATGTGTCTGCTGCCGTGGCCAAGACAGTGCGGATGACACTCGTGATCGTGGTGGTGTACGTGCTGTGCTGGGCCCCATTCTTTCTGGTGCAACTGTGGGCCGCCTGGGATCCTGAAGCTCCTCTTGAAGGCGCTCCCTTCGTGCTGCTGATGCTGCTGGCTAGCCTGAACTCCTGCACAAACCCCTGGATCTACGCCTCCTTCAGCAGCAGCGTGTCCAGCGAGCTGAGAAGCCTGCTGTGTTGTGCCAGAGGCAGGACACCTCCATCTCTGGGACCTCAGGATGAGAGCTGTACCACCGCCTCTTCTAGCCTGGCCAAGGATACAAGTTCTGGCGGCGGAGGATCTGGCGGAGGTGGAAGCGGAGGCGGCGGATCTGAGAGCCTGTTTAAGGGCCCCAGAGACTACAACCCCATCAGCTCCACCATCTGCCACCTGACCAATGAGTCCGATGGCCACACCGCCAGCCTGTACGGCATCGGCTTCGGCCCCTTCATCATCACCAACAAGCACCTGTTCAGACGGAACAACGGCACCCTGCTGGTGCAGTCTCTGCACGGCGTGTTCAAAGTGAAGAATACCACCACACTGCAGCAGCACCTGATCGACGGCCGGGACATGATCATCATCAGAATGCCCAAGGACTTCCCGCCTTTTCCACAGAAGCTGAAGTTCAGAGAGCCCCAGCGCGAGGAAAGAATCTGCCTGGTCACCACCAACTTCCAGACCAAGAGCATGTCCTCCATGGTGTCCGATACCAGCTGCACATTCCCTAGCAGCGACGGCATCTTCTGGAAGCACTGGATTCAGACCAAGGACGGCCAGTGTGGCAGCCCTCTGGTGTCTACAAGAGATGGCTTCATCGTGGGCATCCACAGCGCCAGCAACTTCACCAATACCAACAACTACTTCACCAGCGTGCCGAAGAACTTCATGGAACTGCTGACCAATCAAGAGGCTCAGCAGTGGGTTTCCGGCTGGCGGCTGAATGCTGATTCTGTGCTGTGGGGCGGACACAAGGTGTTCATGGTCTGAATCTAGACCTTCTGCGGGGCTTGCCTTCTGGCCATGCCCTTCTTCTCTCCCTTGCACCTGTACCTCTTGGTCTTTGAATAAAGCCTGAGTAGGAAAAAAAAAAAAAAAAAAAAAAAAAAAAAAAAAAAAAAAAAAAAAAAAAAAAAAAAAAAAAAAAAAAAAAAAAAAAAAAAAAAAAAAAAAAAAAAAAAAAAAAAAAAAAAAAAAAAAAAA]

- **AVPR2-TEVpΔS135N**

T7 promoter (for CleanCap AG Reagent): 11-30

Kozak sequence (including start codon): 79-87

Human AVPR2: 88-1197

3xGGGGS flexible linker: 1198-1242

Carboxy-terminally truncated TEVp (S135N/S219V mutant): 1243-1896

Stop codon: 1897-1899

[CAGTGAATTGTAATACGACTCACTATAAGGCGAATTAAGAGAGAAAAGAAGAGTAAGAAGAAATATAAGACACCGGTCGCCACCATGCTGATGGCCTCTACAACATCTGCCGTGCCTGGACACCCTAGCCTGCCTTCTCTGCCTAGCAACAGCAGCCAAGAGAGGCCCCTGGATACCAGAGATCCTCTGCTGGCCAGAGCCGAACTGGCCCTGCTGTCTATCGTGTTTGTGGCCGTGGCTCTGTCCAACGGACTGGTTCTTGCTGCCCTGGCTCGGAGAGGAAGAAGAGGACATTGGGCCCCTATCCACGTGTTCATCGGCCATCTGTGTCTGGCCGATCTGGCTGTGGCACTGTTTCAGGTTCTGCCTCAGCTGGCCTGGAAGGCCACCGATAGATTCAGAGGCCCCGACGCTCTGTGCAGAGCCGTGAAATACCTGCAGATGGTCGGGATGTACGCCAGCAGCTACATGATCCTGGCCATGACACTGGACCGGCACAGAGCCATCTGTAGACCCATGCTGGCCTACAGACATGGCTCTGGCGCCCACTGGAATAGACCTGTGCTTGTGGCCTGGGCCTTCAGCCTGCTTCTGTCTCTGCCCCAGCTGTTCATCTTCGCCCAGAGAAATGTGGAAGGCGGCAGCGGCGTTACAGATTGCTGGGCCTGTTTTGCCGAGCCTTGGGGTAGAAGAACCTACGTGACATGGATCGCCCTGATGGTGTTCGTGGCCCCTACACTGGGAATTGCCGCTTGCCAGGTGCTGATCTTCAGAGAGATCCACGCCAGCCTGGTGCCAGGACCTTCTGAAAGACCTGGCGGACGCAGACGGGGAAGAAGAACAGGATCTCCTGGCGAAGGCGCCCATGTGTCTGCTGCCGTGGCCAAGACAGTGCGGATGACACTCGTGATCGTGGTGGTGTACGTGCTGTGCTGGGCCCCATTCTTTCTGGTGCAACTGTGGGCCGCCTGGGATCCTGAAGCTCCTCTTGAAGGCGCTCCCTTCGTGCTGCTGATGCTGCTGGCTAGCCTGAACTCCTGCACAAACCCCTGGATCTACGCCTCCTTCAGCAGCAGCGTGTCCAGCGAGCTGAGAAGCCTGCTGTGTTGTGCCAGAGGCAGGACACCTCCATCTCTGGGACCTCAGGATGAGAGCTGTACCACCGCCTCTTCTAGCCTGGCCAAGGATACAAGTTCTGGCGGCGGAGGATCTGGCGGAGGTGGAAGCGGAGGCGGCGGATCTGAGAGCCTGTTTAAGGGCCCCAGAGACTACAACCCCATCAGCTCCACCATCTGCCACCTGACCAATGAGTCCGATGGCCACACCACAAGCCTGTACGGCATCGGCTTCGGCCCCTTCATCATCACCAACAAGCACCTGTTCAGACGGAACAACGGCACCCTGCTGGTGCAGTCTCTGCACGGCGTGTTCAAAGTGAAGAATACCACCACACTGCAGCAGCACCTGATCGACGGCCGGGACATGATCATCATCAGAATGCCCAAGGACTTCCCGCCTTTTCCACAGAAGCTGAAGTTCAGAGAGCCCCAGCGCGAGGAAAGAATCTGCCTGGTCACCACCAACTTCCAGACCAAGAGCATGTCCTCCATGGTGTCCGATACCAGCTGCACATTCCCTAGCAGCGACGGCATCTTCTGGAAGCACTGGATTCAGACCAAGGACGGCCAGTGTGGCAACCCTCTGGTGTCTACAAGAGATGGCTTCATCGTGGGCATCCACAGCGCCAGCAACTTCACCAATACCAACAACTACTTCACCAGCGTGCCGAAGAACTTCATGGAACTGCTGACCAATCAAGAGGCTCAGCAGTGGGTTTCCGGCTGGCGGCTGAATGCTGATTCTGTGCTGTGGGGCGGACACAAGGTGTTCATGGTCTGAATCTAGACCTTCTGCGGGGCTTGCCTTCTGGCCATGCCCTTCTTCTCTCCCTTGCACCTGTACCTCTTGGTCTTTGAATAAAGCCTGAGTAGGAAAAAAAAAAAAAAAAAAAAAAAAAAAAAAAAAAAAAAAAAAAAAAAAAAAAAAAAAAAAAAAAAAAAAAAAAAAAAAAAAAAAAAAAAAAAAAAAAAAAAAAAAAAAAAAAAAAAAAAA]

- **EP3-TEVpΔT30A**

T7 promoter (for CleanCap AG Reagent): 11-30

Kozak sequence (including start codon): 79-87

Human EP3 isoform I: 88-1254

A343-S371 of human AVPR2: 1255-1341

3xGGGGS flexible linker: 1342-1386

Carboxy-terminally truncated TEVp (T30A/S219V mutant): 1387-2040

Stop codon: 2041-2043

[CAGTGAATTGTAATACGACTCACTATAAGGCGAATTAAGAGAGAAAAGAAGAGTAAGAAGAAATATAAGACACCGGTCGCCACCATGAAGGAGACCCGGGGCTACGGCGGCGATGCCCCCTTCTGCACCAGGCTGAACCACAGCTACACCGGCATGTGGGCCCCCGAGCGGAGTGCTGAGGCCAGGGGAAACCTGACCCGGCCTCCTGGCAGCGGAGAGGACTGCGGCAGCGTGAGTGTGGCTTTCCCTATCACCATGCTGCTGACCGGCTTCGTGGGCAACGCTCTTGCCATGCTGCTGGTGAGCCGGAGCTACCGGCGGAGGGAGAGCAAGCGGAAGAAGAGCTTCCTGCTGTGCATCGGCTGGCTGGCCCTGACCGACCTGGTGGGCCAGCTGCTGACCACCCCCGTGGTGATCGTGGTGTACCTGAGCAAGCAGCGGTGGGAGCACATCGACCCCAGCGGCCGGCTGTGCACCTTCTTCGGCCTGACCATGACCGTGTTCGGCCTGAGCAGCCTGTTCATCGCCAGCGCCATGGCCGTGGAGAGGGCCCTTGCCATCAGGGCTCCCCACTGGTACGCCAGCCACATGAAGACCCGGGCCACCCGGGCTGTTCTGCTGGGAGTGTGGCTTGCCGTGCTGGCCTTCGCTCTGCTGCCTGTGCTGGGCGTGGGCCAGTACACCGTGCAATGGCCTGGCACATGGTGCTTCATCAGCACCGGCAGGGGAGGCAACGGCACCAGCAGCAGCCACAACTGGGGCAACCTGTTCTTCGCCAGTGCCTTCGCCTTCCTGGGCCTGCTGGCCCTGACCGTGACCTTCAGCTGCAACCTGGCCACCATCAAGGCCCTGGTGAGCCGGTGCCGGGCCAAGGCCACCGCTAGTCAGAGCAGCGCCCAGTGGGGACGGATCACCACAGAGACCGCCATCCAGCTGATGGGCATCATGTGCGTGCTGAGCGTGTGCTGGAGCCCCCTGCTGATCATGATGCTGAAGATGATCTTCAACCAGACCAGCGTGGAGCACTGCAAGACCCACACCGAGAAGCAGAAGGAGTGCAACTTCTTCCTGATCGCCGTGCGGCTGGCCAGCCTGAACCAGATCCTGGACCCCTGGGTGTACCTGCTGCTGCGGAAGATCCTGCTGCGGAAGTTCTGCCAGATCCGGTACCACACCAACAACTACGCCAGCAGCAGCACCAGCCTGCCCTGCCAGTGCAGCAGCACCCTGATGTGGAGCGACCACCTGGAGCGGGCCAGAGGCAGGACACCTCCATCTCTGGGACCTCAGGATGAGAGCTGTACCACCGCCTCTTCTAGCCTGGCCAAGGATACAAGTTCTGGCGGCGGAGGATCTGGCGGAGGTGGAAGCGGAGGCGGCGGATCTGAGAGCCTGTTTAAGGGCCCCAGAGACTACAACCCCATCAGCTCCACCATCTGCCACCTGACCAATGAGTCCGATGGCCACACCGCCAGCCTGTACGGCATCGGCTTCGGCCCCTTCATCATCACCAACAAGCACCTGTTCAGACGGAACAACGGCACCCTGCTGGTGCAGTCTCTGCACGGCGTGTTCAAAGTGAAGAATACCACCACACTGCAGCAGCACCTGATCGACGGCCGGGACATGATCATCATCAGAATGCCCAAGGACTTCCCGCCTTTTCCACAGAAGCTGAAGTTCAGAGAGCCCCAGCGCGAGGAAAGAATCTGCCTGGTCACCACCAACTTCCAGACCAAGAGCATGTCCTCCATGGTGTCCGATACCAGCTGCACATTCCCTAGCAGCGACGGCATCTTCTGGAAGCACTGGATTCAGACCAAGGACGGCCAGTGTGGCAGCCCTCTGGTGTCTACAAGAGATGGCTTCATCGTGGGCATCCACAGCGCCAGCAACTTCACCAATACCAACAACTACTTCACCAGCGTGCCGAAGAACTTCATGGAACTGCTGACCAATCAAGAGGCTCAGCAGTGGGTTTCCGGCTGGCGGCTGAATGCTGATTCTGTGCTGTGGGGCGGACACAAGGTGTTCATGGTCTGAATCTAGACCTTCTGCGGGGCTTGCCTTCTGGCCATGCCCTTCTTCTCTCCCTTGCACCTGTACCTCTTGGTCTTTGAATAAAGCCTGAGTAGGAAAAAAAAAAAAAAAAAAAAAAAAAAAAAAAAAAAAAAAAAAAAAAAAAAAAAAAAAAAAAAAAAAAAAAAAAAAAAAAAAAAAAAAAAAAAAAAAAAAAAAAAAAAAAAAAAAAAAAAA]

- **EP4-TEVpΔT30A**

T7 promoter (for CleanCap AG Reagent): 11-30

Kozak sequence (including start codon): 79-87

Human EP4: 88-1548

A343-S371 of AVPR2: 1549-1635

3xGGGGS flexible linker: 1636-1680

Carboxy-terminally truncated TEVp (T30A/S219V mutant): 1681-2334

Stop codon: 2335-2337

[CAGTGAATTGTAATACGACTCACTATAAGGCGAATTAAGAGAGAAAAGAAGAGTAAGAAGAAATATAAGACACCGGTCGCCACCATGAGCACCCCCGGCGTGAACAGCAGCGCCAGCCTGAGCCCCGACAGGCTGAACAGCCCCGTGACCATCCCCGCCGTGATGTTCATCTTCGGCGTGGTGGGCAACCTGGTGGCCATCGTGGTGCTGTGCAAGAGCCGGAAGGAGCAGAAGGAGACCACCTTCTACACCCTGGTGTGCGGCCTGGCCGTGACCGACCTGCTGGGCACCCTGCTGGTGAGCCCCGTGACCATCGCTACCTACATGAAGGGCCAGTGGCCCGGCGGACAGCCTCTGTGCGAGTACAGCACCTTCATCCTGCTGTTCTTCAGCCTGAGCGGCCTGAGCATCATCTGCGCCATGAGCGTGGAGCGGTACCTGGCCATCAACCACGCCTACTTCTACAGCCACTACGTGGACAAGCGGCTGGCCGGCCTGACCCTGTTCGCCGTGTACGCCAGCAACGTGCTGTTCTGCGCCCTGCCCAACATGGGCCTGGGCAGCAGCCGGCTGCAGTACCCTGACACCTGGTGCTTCATCGACTGGACCACCAACGTGACCGCCCACGCCGCCTACAGCTACATGTACGCCGGCTTCAGCAGCTTCCTGATCCTGGCCACCGTGCTGTGCAACGTGCTGGTGTGCGGCGCCCTGCTGCGGATGCACCGGCAGTTCATGCGGCGGACCAGCCTGGGCACCGAGCAACACCATGCCGCTGCTGCTGCCAGCGTTGCTAGTCGGGGACACCCTGCTGCCAGTCCCGCCCTGCCTAGGCTGAGCGATTTTCGGAGGCGGAGGAGCTTCAGGAGGATTGCCGGAGCCGAGATCCAGATGGTGATCCTGCTGATCGCCACCAGCCTGGTGGTGCTGATCTGCAGCATCCCCCTGGTGGTGCGGGTGTTCGTGAACCAGCTGTACCAGCCCAGCCTGGAGCGGGAGGTGAGCAAGAACCCCGACCTGCAGGCCATCCGGATCGCCAGCGTGAACCCCATCCTGGACCCCTGGATCTACATCCTGCTGCGGAAGACCGTGCTGAGCAAGGCCATCGAGAAGATCAAGTGCCTGTTCTGCCGGATCGGCGGCAGCCGGAGGGAGAGGAGCGGCCAACACTGCAGCGACAGCCAGCGGACCAGCAGCGCCATGAGCGGCCACAGCAGGAGTTTCATCAGCCGGGAGCTGAAGGAGATCAGCAGCACCAGCCAGACCCTGCTGCCCGACCTGAGCCTGCCCGACCTGAGCGAGAACGGCCTGGGAGGAAGGAACCTGCTTCCCGGAGTGCCCGGCATGGGCCTGGCTCAGGAGGACACCACCAGCCTGCGGACCCTGCGGATCAGCGAGACCAGCGACAGCAGCCAGGGCCAAGATAGTGAGAGCGTGCTGCTGGTTGATGAGGCTGGAGGAAGCGGCAGGGCCGGACCTGCTCCCAAGGGCAGCAGCCTGCAGGTGACCTTCCCCAGCGAGACCCTGAACCTGAGCGAGAAGTGCATCGCCAGAGGCAGGACACCTCCATCTCTGGGACCTCAGGATGAGAGCTGTACCACCGCCTCTTCTAGCCTGGCCAAGGATACAAGTTCTGGCGGCGGAGGATCTGGCGGAGGTGGAAGCGGAGGCGGCGGATCTGAGAGCCTGTTTAAGGGCCCCAGAGACTACAACCCCATCAGCTCCACCATCTGCCACCTGACCAATGAGTCCGATGGCCACACCGCCAGCCTGTACGGCATCGGCTTCGGCCCCTTCATCATCACCAACAAGCACCTGTTCAGACGGAACAACGGCACCCTGCTGGTGCAGTCTCTGCACGGCGTGTTCAAAGTGAAGAATACCACCACACTGCAGCAGCACCTGATCGACGGCCGGGACATGATCATCATCAGAATGCCCAAGGACTTCCCGCCTTTTCCACAGAAGCTGAAGTTCAGAGAGCCCCAGCGCGAGGAAAGAATCTGCCTGGTCACCACCAACTTCCAGACCAAGAGCATGTCCTCCATGGTGTCCGATACCAGCTGCACATTCCCTAGCAGCGACGGCATCTTCTGGAAGCACTGGATTCAGACCAAGGACGGCCAGTGTGGCAGCCCTCTGGTGTCTACAAGAGATGGCTTCATCGTGGGCATCCACAGCGCCAGCAACTTCACCAATACCAACAACTACTTCACCAGCGTGCCGAAGAACTTCATGGAACTGCTGACCAATCAAGAGGCTCAGCAGTGGGTTTCCGGCTGGCGGCTGAATGCTGATTCTGTGCTGTGGGGCGGACACAAGGTGTTCATGGTCTGAATCTAGACCTTCTGCGGGGCTTGCCTTCTGGCCATGCCCTTCTTCTCTCCCTTGCACCTGTACCTCTTGGTCTTTGAATAAAGCCTGAGTAGGAAAAAAAAAAAAAAAAAAAAAAAAAAAAAAAAAAAAAAAAAAAAAAAAAAAAAAAAAAAAAAAAAAAAAAAAAAAAAAAAAAAAAAAAAAAAAAAAAAAAAAAAAAAAAAAAAAAAAAAA]

- **B2-TEVpΔT30A**

T7 promoter (for CleanCap AG Reagent): 11-30

Kozak sequence (including start codon): 79-87

Human B2: 88-1257

A343-S371 of human AVPR2: 1258-1344

3xGGGGS flexible linker: 1345-1389

Carboxy-terminally truncated TEVp (T30A/S219V mutant): 1390-2043

Stop codon: 2044-2046

[CAGTGAATTGTAATACGACTCACTATAAGGCGAATTAAGAGAGAAAAGAAGAGTAAGAAGAAATATAAGACACCGGTCGCCACCATGTTTAGCCCTTGGAAGATCAGCATGTTCCTGAGCGTGCGCGAGGATAGCGTGCCAACCACAGCCAGCTTTAGCGCCGACATGCTGAACGTGACACTGCAGGGCCCTACACTGAACGGCACATTCGCCCAGAGCAAGTGCCCTCAGGTGGAATGGCTCGGCTGGCTGAATACGATCCAGCCTCCTTTCCTGTGGGTGCTGTTCGTGCTGGCCACACTGGAAAACATCTTCGTGCTGAGCGTGTTCTGCCTGCACAAGAGCAGCTGTACCGTGGCCGAGATCTACCTGGGAAATCTGGCCGCTGCCGATCTGATCCTGGCTTGCGGACTTCCTTTCTGGGCCATCACCATCAGCAACAACTTCGACTGGCTGTTCGGCGAGACACTGTGCAGAGTGGTCAACGCCATCATCAGCATGAACCTGTACAGCAGCATCTGCTTTCTGATGCTGGTGTCCATCGACCGGTATCTGGCCCTGGTCAAGACCATGAGCATGGGCAGAATGCGGGGCGTCAGATGGGCCAAGCTGTACTCTCTGGTCATCTGGGGCTGTACCCTGCTGCTGTCTAGCCCCATGCTCGTGTTCCGGACCATGAAGGAATACTCCGACGAGGGCCACAACGTGACCGCCTGTGTGATCAGCTACCCCAGCCTGATCTGGGAAGTGTTCACCAACATGCTGCTGAATGTCGTGGGCTTCCTGCTGCCTCTGAGCGTGATCACCTTCTGCACCATGCAGATCATGCAGGTCCTGCGGAACAACGAGATGCAGAAGTTCAAAGAGATCCAGACCGAGCGGAGAGCCACCGTGCTGGTTCTGGTTGTGCTGCTCCTGTTCATCATCTGCTGGCTGCCCTTCCAGATCAGCACCTTCCTGGACACCCTGCACAGACTGGGCATCCTGTCCAGCTGCCAGGACGAGAGAATCATCGATGTGATCACCCAGATCGCCAGCTTCATGGCCTACAGCAACAGCTGCCTGAATCCTCTGGTGTACGTGATCGTGGGCAAGCGCTTCAGAAAGAAAAGCTGGGAAGTCTACCAGGGCGTGTGCCAGAAAGGCGGCTGTAGATCTGAGCCCATCCAGATGGAAAACTCCATGGGCACCCTGCGGACCAGCATCTCCGTGGAAAGACAGATCCACAAGCTGCAGGATTGGGCCGGCTCTAGACAAGCCAGAGGCAGGACACCTCCATCTCTGGGACCTCAGGATGAGAGCTGTACCACCGCCTCTTCTAGCCTGGCCAAGGATACAAGTTCTGGCGGCGGAGGATCTGGCGGAGGTGGAAGCGGAGGCGGCGGATCTGAGAGCCTGTTTAAGGGCCCCAGAGACTACAACCCCATCAGCTCCACCATCTGCCACCTGACCAATGAGTCCGATGGCCACACCGCCAGCCTGTACGGCATCGGCTTCGGCCCCTTCATCATCACCAACAAGCACCTGTTCAGACGGAACAACGGCACCCTGCTGGTGCAGTCTCTGCACGGCGTGTTCAAAGTGAAGAATACCACCACACTGCAGCAGCACCTGATCGACGGCCGGGACATGATCATCATCAGAATGCCCAAGGACTTCCCGCCTTTTCCACAGAAGCTGAAGTTCAGAGAGCCCCAGCGCGAGGAAAGAATCTGCCTGGTCACCACCAACTTCCAGACCAAGAGCATGTCCTCCATGGTGTCCGATACCAGCTGCACATTCCCTAGCAGCGACGGCATCTTCTGGAAGCACTGGATTCAGACCAAGGACGGCCAGTGTGGCAGCCCTCTGGTGTCTACAAGAGATGGCTTCATCGTGGGCATCCACAGCGCCAGCAACTTCACCAATACCAACAACTACTTCACCAGCGTGCCGAAGAACTTCATGGAACTGCTGACCAATCAAGAGGCTCAGCAGTGGGTTTCCGGCTGGCGGCTGAATGCTGATTCTGTGCTGTGGGGCGGACACAAGGTGTTCATGGTCTGAATCTAGACCTTCTGCGGGGCTTGCCTTCTGGCCATGCCCTTCTTCTCTCCCTTGCACCTGTACCTCTTGGTCTTTGAATAAAGCCTGAGTAGGAAAAAAAAAAAAAAAAAAAAAAAAAAAAAAAAAAAAAAAAAAAAAAAAAAAAAAAAAAAAAAAAAAAAAAAAAAAAAAAAAAAAAAAAAAAAAAAAAAAAAAAAAAAAAAAAAAAAAAAA]

- **EP3V2-TEVpΔT30A**

T7 promoter (for CleanCap AG Reagent): 11-30

Kozak sequence (including start codon): 79-87

K2-R350 of human EP3 isoform I: 88-1134

S327-S371 of human AVPR2: 1135-1269

3xGGGGS flexible linker: 1270-1314

Carboxy-terminally truncated TEVp (T30A/S219V mutant): 1315-1968

Stop codon: 1969-1971

[CAGTGAATTGTAATACGACTCACTATAAGGCGAATTAAGAGAGAAAAGAAGAGTAAGAAGAAATATAAGACACCGGTCGCCACCATGAAGGAGACCCGGGGCTACGGCGGCGATGCCCCCTTCTGCACCAGGCTGAACCACAGCTACACCGGCATGTGGGCCCCCGAGCGGAGTGCTGAGGCCAGGGGAAACCTGACCCGGCCTCCTGGCAGCGGAGAGGACTGCGGCAGCGTGAGTGTGGCTTTCCCTATCACCATGCTGCTGACCGGCTTCGTGGGCAACGCTCTTGCCATGCTGCTGGTGAGCCGGAGCTACCGGCGGAGGGAGAGCAAGCGGAAGAAGAGCTTCCTGCTGTGCATCGGCTGGCTGGCCCTGACCGACCTGGTGGGCCAGCTGCTGACCACCCCCGTGGTGATCGTGGTGTACCTGAGCAAGCAGCGGTGGGAGCACATCGACCCCAGCGGCCGGCTGTGCACCTTCTTCGGCCTGACCATGACCGTGTTCGGCCTGAGCAGCCTGTTCATCGCCAGCGCCATGGCCGTGGAGAGGGCCCTTGCCATCAGGGCTCCCCACTGGTACGCCAGCCACATGAAGACCCGGGCCACCCGGGCTGTTCTGCTGGGAGTGTGGCTTGCCGTGCTGGCCTTCGCTCTGCTGCCTGTGCTGGGCGTGGGCCAGTACACCGTGCAATGGCCTGGCACATGGTGCTTCATCAGCACCGGCAGGGGAGGCAACGGCACCAGCAGCAGCCACAACTGGGGCAACCTGTTCTTCGCCAGTGCCTTCGCCTTCCTGGGCCTGCTGGCCCTGACCGTGACCTTCAGCTGCAACCTGGCCACCATCAAGGCCCTGGTGAGCCGGTGCCGGGCCAAGGCCACCGCTAGTCAGAGCAGCGCCCAGTGGGGACGGATCACCACAGAGACCGCCATCCAGCTGATGGGCATCATGTGCGTGCTGAGCGTGTGCTGGAGCCCCCTGCTGATCATGATGCTGAAGATGATCTTCAACCAGACCAGCGTGGAGCACTGCAAGACCCACACCGAGAAGCAGAAGGAGTGCAACTTCTTCCTGATCGCCGTGCGGCTGGCCAGCCTGAACCAGATCCTGGACCCCTGGGTGTACCTGCTGCTGCGGTCCTTCAGCAGCAGCGTGTCCAGCGAGCTGAGAAGCCTGCTGTGTTGTGCCAGAGGCAGGACACCTCCATCTCTGGGACCTCAGGATGAGAGCTGTACCACCGCCTCTTCTAGCCTGGCCAAGGATACAAGTTCTGGCGGCGGAGGATCTGGCGGAGGTGGAAGCGGAGGCGGCGGATCTGAGAGCCTGTTTAAGGGCCCCAGAGACTACAACCCCATCAGCTCCACCATCTGCCACCTGACCAATGAGTCCGATGGCCACACCGCCAGCCTGTACGGCATCGGCTTCGGCCCCTTCATCATCACCAACAAGCACCTGTTCAGACGGAACAACGGCACCCTGCTGGTGCAGTCTCTGCACGGCGTGTTCAAAGTGAAGAATACCACCACACTGCAGCAGCACCTGATCGACGGCCGGGACATGATCATCATCAGAATGCCCAAGGACTTCCCGCCTTTTCCACAGAAGCTGAAGTTCAGAGAGCCCCAGCGCGAGGAAAGAATCTGCCTGGTCACCACCAACTTCCAGACCAAGAGCATGTCCTCCATGGTGTCCGATACCAGCTGCACATTCCCTAGCAGCGACGGCATCTTCTGGAAGCACTGGATTCAGACCAAGGACGGCCAGTGTGGCAGCCCTCTGGTGTCTACAAGAGATGGCTTCATCGTGGGCATCCACAGCGCCAGCAACTTCACCAATACCAACAACTACTTCACCAGCGTGCCGAAGAACTTCATGGAACTGCTGACCAATCAAGAGGCTCAGCAGTGGGTTTCCGGCTGGCGGCTGAATGCTGATTCTGTGCTGTGGGGCGGACACAAGGTGTTCATGGTCTGAATCTAGACCTTCTGCGGGGCTTGCCTTCTGGCCATGCCCTTCTTCTCTCCCTTGCACCTGTACCTCTTGGTCTTTGAATAAAGCCTGAGTAGGAAAAAAAAAAAAAAAAAAAAAAAAAAAAAAAAAAAAAAAAAAAAAAAAAAAAAAAAAAAAAAAAAAAAAAAAAAAAAAAAAAAAAAAAAAAAAAAAAAAAAAAAAAAAAAAAAAAAAAAA]

- **2xScKt-Luc2**

T7 promoter (for CleanCap AG Reagent): 11-30

Stabilized 2xKink-turn: 31-99

Kozak sequence (including start codon): 120-128

Luc2: 129-1775

Stop codon: 1776-1778

[CAGTGAATTGTAATACGACTCACTATAAGGGAGAGGGGTGAGCGGGCCCGCGATGATCCCCTCGACTTCTGACGCACGTGCGGATGATGAAGTCTTCCCAGAAATATAAGACACCGGTCGCCACCATGGAAGATGCCAAAAACATTAAGAAGGGCCCAGCGCCATTCTACCCACTCGAAGACGGGACCGCCGGCGAGCAGCTGCACAAAGCCATGAAGCGCTACGCCCTGGTGCCCGGCACCATCGCCTTTACCGACGCACATATCGAGGTGGACATTACCTACGCCGAGTACTTCGAGATGAGCGTTCGGCTGGCAGAAGCTATGAAGCGCTATGGGCTGAATACAAACCATCGGATCGTGGTGTGCAGCGAGAATAGCTTGCAGTTCTTCATGCCCGTGTTGGGTGCCCTGTTCATCGGTGTGGCTGTGGCCCCAGCTAACGACATCTACAACGAGCGCGAGCTGCTGAACAGCATGGGCATCAGCCAGCCCACCGTCGTATTCGTGAGCAAGAAAGGGCTGCAAAAGATCCTCAACGTGCAAAAGAAGCTACCGATCATACAAAAGATCATCATCATGGATAGCAAGACCGACTACCAGGGCTTCCAAAGCATGTACACCTTCGTGACTTCCCATTTGCCACCCGGCTTCAACGAGTACGACTTCGTGCCCGAGAGCTTCGACCGGGACAAAACCATCGCCCTGATCATGAACAGTAGTGGCAGTACCGGATTGCCCAAGGGCGTAGCCCTACCGCACCGCACCGCTTGTGTCCGATTCAGTCATGCCCGCGACCCCATCTTCGGCAACCAGATCATCCCCGACACCGCTATCCTCAGCGTGGTGCCATTTCACCACGGCTTCGGCATGTTCACCACGCTGGGCTACTTGATCTGCGGCTTTCGGGTCGTGCTCATGTACCGCTTCGAGGAGGAGCTATTCTTGCGCAGCTTGCAAGACTATAAGATTCAATCTGCCCTGCTGGTGCCCACACTATTTAGCTTCTTCGCTAAGAGCACTCTCATCGACAAGTACGACCTAAGCAACTTGCACGAGATCGCCAGCGGCGGGGCGCCGCTCAGCAAGGAGGTAGGTGAGGCCGTGGCCAAACGCTTCCACCTACCAGGCATCCGCCAGGGCTACGGCCTGACAGAAACAACCAGCGCCATTCTGATCACCCCCGAAGGGGACGACAAGCCTGGCGCAGTAGGCAAGGTGGTGCCCTTCTTCGAGGCTAAGGTGGTGGACTTGGACACCGGTAAGACACTGGGTGTGAACCAGCGCGGCGAGCTGTGCGTCCGTGGCCCCATGATCATGAGCGGCTACGTTAACAACCCCGAGGCTACAAACGCTCTCATCGACAAGGACGGCTGGCTGCACAGCGGCGACATCGCCTACTGGGACGAGGACGAGCACTTCTTCATCGTGGACCGGCTGAAGAGCCTGATCAAATACAAGGGCTACCAGGTAGCCCCAGCCGAACTGGAGAGCATCCTGCTGCAACACCCCAACATCTTCGACGCCGGGGTCGCCGGCCTGCCCGACGACGATGCCGGCGAGCTGCCCGCCGCAGTCGTCGTGCTGGAACACGGTAAAACCATGACCGAGAAGGAGATCGTGGACTATGTGGCCAGCCAGGTTACAACCGCCAAGAAGCTGCGCGGTGGTGTTGTGTTCGTGGACGAGGTGCCTAAAGGACTGACCGGCAAGTTGGACGCCCGCAAGATCCGCGAGATTCTCATTAAGGCCAAGAAGGGCGGCAAGATCGCCGTGTAAATCTAGACCTTCTGCGGGGCTTGCCTTCTGGCCATGCCCTTCTTCTCTCCCTTGCACCTGTACCTCTTGGTCTTTGAATAAAGCCTGAGTAGGAAAAAAAAAAAAAAAAAAAAAAAAAAAAAAAAAAAAAAAAAAAAAAAAAAAAAAAAAAAAAAAAAAAAAAAAAAAAAAAAAAAAAAAAAAAAAAAAAAAAAAAAAAAAAAAAAAAAAAAA]

- **ARRB2-tevD-L7Ae**

T7 promoter (for CleanCap AG Reagent): 11-30

Kozak sequence (including start codon): 79-87

ARRB2: 88-1311

3xGGGGS flexible linker: 1312-1356

tevD: 1357-1419

2xGGGGS flexible linker: 1420-1449

L7Ae: 1450-1803

Stop codon: 1804-1806

[CAGTGAATTGTAATACGACTCACTATAAGGCGAATTAAGAGAGAAAAGAAGAGTAAGAAGAAATATAAGACACCGGTCGCCACCATGGGAGAAAAGCCTGGCACCAGAGTGTTCAAGAAGTCTAGCCCCAACTGCAAGCTGACCGTGTACCTCGGCAAGCGGGACTTCGTGGACCACCTGGACAAAGTGGATCCCGTGGATGGCGTGGTGCTGGTGGATCCTGACTACCTGAAGGACCGGAAGGTGTTCGTGACCCTGACCTGCGCCTTCAGATACGGCAGAGAAGATCTGGACGTGCTGGGCCTGAGCTTCAGAAAGGACCTGTTTATCGCCACCTACCAGGCCTTTCCACCTGTGCCTAATCCTCCACGGCCTCCTACCAGACTGCAGGACAGACTGCTGAGAAAGCTGGGCCAGCACGCTCACCCCTTTTTCTTCACAATCCCTCAAAACCTGCCTTGCAGCGTGACACTGCAGCCCGGACCTGAGGATACAGGCAAAGCTTGCGGCGTGGACTTCGAGATCAGAGCCTTCTGTGCCAAGAGCCTGGAAGAGAAGTCCCACAAGAGAAACAGCGTGCGGCTGGTCATCAGAAAGGTGCAGTTCGCCCCTGAGAAGCCTGGACCTCAGCCTTCTGCCGAGACAACCAGACACTTCCTGATGAGCGACCGCAGCCTGCATCTGGAAGCCAGCCTCGACAAAGAGCTGTACTACCACGGCGAGCCCCTGAACGTCAACGTCCACGTGACCAACAACAGCACCAAGACCGTGAAAAAGATCAAGGTGTCCGTGCGGCAGTACGCCGACATCTGCCTGTTTAGCACAGCCCAGTACAAGTGCCCCGTGGCTCAGCTGGAACAGGACGATCAAGTGTCCCCTAGCAGCACCTTCTGCAAGGTGTACACAATCACCCCTCTGCTGAGCGACAACAGAGAGAAGAGAGGACTGGCCCTGGACGGCAAGCTGAAGCACGAGGATACAAACCTGGCCAGCAGCACCATCGTGAAAGAGGGCGCCAACAAAGAGGTGCTGGGCATCCTGGTGTCCTACAGAGTGAAAGTGAAGCTGGTGGTGTCCAGAGGCGGCGACGTTTCAGTGGAACTGCCCTTCGTGCTGATGCACCCCAAGCCTCACGATCACATCCCTCTGCCTAGACCTCAGAGCGCCGCTCCTGAAACAGATGTGCCCGTGGACACCAACCTGATCGAGTTCGACACCAACTACGCCACCGACGACGACATCGTGTTCGAGGACTTCGCCCGGCTGAGACTGAAGGGCATGAAGGACGACGACTACGACGATCAGCTTTGTGGCGGCGGAGGATCTGGCGGAGGTGGAAGCGGAGGCGGTGGATCTGAGAACCTGTACTTCCAGTTCCACAAGAGCGGCGCCTGGAAGCTGCCTGTGTCTCTGGTTAAGGGCGGCGGCGGCTCCGGCGGCGGCGGCTCCTACGTGCGGTTCGAGGTGCCCGAGGACATGCAGAACGAGGCCCTGAGCCTGCTGGAGAAGGTGCGGGAGAGCGGCAAGGTGAAGAAGGGCACCAACGAGACCACCAAGGCCGTGGAGCGGGGCCTTGCCAAGCTGGTGTACATCGCCGAGGACGTGGACCCCCCCGAGATCGTGGCCCACCTTCCTCTGCTGTGCGAGGAGAAGAACGTGCCCTACATCTACGTGAAGAGCAAGAACGACCTGGGCCGGGCCGTTGGCATCGAGGTGCCCTGCGCCAGTGCTGCCATCATCAACGAGGGCGAGCTGCGGAAGGAGCTGGGCAGCCTGGTGGAGAAGATCAAGGGCCTGCAGAAGTGAATCTAGACCTTCTGCGGGGCTTGCCTTCTGGCCATGCCCTTCTTCTCTCCCTTGCACCTGTACCTCTTGGTCTTTGAATAAAGCCTGAGTAGGAAAAAAAAAAAAAAAAAAAAAAAAAAAAAAAAAAAAAAAAAAAAAAAAAAAAAAAAAAAAAAAAAAAAAAAAAAAAAAAAAAAAAAAAAAAAAAAAAAAAAAAAAAAAAAAAAAAAAAAA]

- **ARRB2-tevD-L7AeCS2**

T7 promoter (for CleanCap AG Reagent): 11-30

Kozak sequence (including start codon): 79-87

ARRB2: 88-1311

3xGGGGS flexible linker: 1312-1356

tevD: 1357-1419

2xGGGGS flexible linker: 1420-1449

L7AeCS2: 1450-1824

TEVp cleavage site: 1615-1635

Stop codon: 1825-1827

[CAGTGAATTGTAATACGACTCACTATAAGGCGAATTAAGAGAGAAAAGAAGAGTAAGAAGAAATATAAGACACCGGTCGCCACCATGGGAGAAAAGCCTGGCACCAGAGTGTTCAAGAAGTCTAGCCCCAACTGCAAGCTGACCGTGTACCTCGGCAAGCGGGACTTCGTGGACCACCTGGACAAAGTGGATCCCGTGGATGGCGTGGTGCTGGTGGATCCTGACTACCTGAAGGACCGGAAGGTGTTCGTGACCCTGACCTGCGCCTTCAGATACGGCAGAGAAGATCTGGACGTGCTGGGCCTGAGCTTCAGAAAGGACCTGTTTATCGCCACCTACCAGGCCTTTCCACCTGTGCCTAATCCTCCACGGCCTCCTACCAGACTGCAGGACAGACTGCTGAGAAAGCTGGGCCAGCACGCTCACCCCTTTTTCTTCACAATCCCTCAAAACCTGCCTTGCAGCGTGACACTGCAGCCCGGACCTGAGGATACAGGCAAAGCTTGCGGCGTGGACTTCGAGATCAGAGCCTTCTGTGCCAAGAGCCTGGAAGAGAAGTCCCACAAGAGAAACAGCGTGCGGCTGGTCATCAGAAAGGTGCAGTTCGCCCCTGAGAAGCCTGGACCTCAGCCTTCTGCCGAGACAACCAGACACTTCCTGATGAGCGACCGCAGCCTGCATCTGGAAGCCAGCCTCGACAAAGAGCTGTACTACCACGGCGAGCCCCTGAACGTCAACGTCCACGTGACCAACAACAGCACCAAGACCGTGAAAAAGATCAAGGTGTCCGTGCGGCAGTACGCCGACATCTGCCTGTTTAGCACAGCCCAGTACAAGTGCCCCGTGGCTCAGCTGGAACAGGACGATCAAGTGTCCCCTAGCAGCACCTTCTGCAAGGTGTACACAATCACCCCTCTGCTGAGCGACAACAGAGAGAAGAGAGGACTGGCCCTGGACGGCAAGCTGAAGCACGAGGATACAAACCTGGCCAGCAGCACCATCGTGAAAGAGGGCGCCAACAAAGAGGTGCTGGGCATCCTGGTGTCCTACAGAGTGAAAGTGAAGCTGGTGGTGTCCAGAGGCGGCGACGTTTCAGTGGAACTGCCCTTCGTGCTGATGCACCCCAAGCCTCACGATCACATCCCTCTGCCTAGACCTCAGAGCGCCGCTCCTGAAACAGATGTGCCCGTGGACACCAACCTGATCGAGTTCGACACCAACTACGCCACCGACGACGACATCGTGTTCGAGGACTTCGCCCGGCTGAGACTGAAGGGCATGAAGGACGACGACTACGACGATCAGCTTTGTGGCGGCGGAGGATCTGGCGGAGGTGGAAGCGGAGGCGGTGGATCTGAGAACCTGTACTTCCAGTTCCACAAGAGCGGCGCCTGGAAGCTGCCTGTGTCTCTGGTTAAGGGCGGCGGCGGCTCCGGCGGCGGCGGCTCCTACGTGCGGTTCGAGGTGCCCGAGGACATGCAGAACGAGGCCCTGAGCCTGCTGGAGAAGGTGCGGGAGAGCGGCAAGGTGAAGAAGGGCACCAACGAGACCACCAAGGCCGTGGAGCGGGGCCTTGCCAAGCTGGTGTACATCGCCGAGGACGTGGACCCCCCCGAGAACCTGTACTTCCAGGGCGAGATCGTGGCCCACCTTCCTCTGCTGTGCGAGGAGAAGAACGTGCCCTACATCTACGTGAAGAGCAAGAACGACCTGGGCCGGGCCGTTGGCATCGAGGTGCCCTGCGCCAGTGCTGCCATCATCAACGAGGGCGAGCTGCGGAAGGAGCTGGGCAGCCTGGTGGAGAAGATCAAGGGCCTGCAGAAGTGAATCTAGACCTTCTGCGGGGCTTGCCTTCTGGCCATGCCCTTCTTCTCTCCCTTGCACCTGTACCTCTTGGTCTTTGAATAAAGCCTGAGTAGGAAAAAAAAAAAAAAAAAAAAAAAAAAAAAAAAAAAAAAAAAAAAAAAAAAAAAAAAAAAAAAAAAAAAAAAAAAAAAAAAAAAAAAAAAAAAAAAAAAAAAAAAAAAAAAAAAAAAAAAA]

- **ARRB2-tevD-L7AeCS3**

T7 promoter (for CleanCap AG Reagent): 11-30

Kozak sequence (including start codon): 79-87

ARRB2: 88-1311

3xGGGGS flexible linker: 1312-1356

tevD: 1357-1419

2xGGGGS flexible linker: 1420-1449

L7AeCS3: 1450-1821

TEVp cleavage site: 1678-1698

Stop codon: 1822-1824

[CAGTGAATTGTAATACGACTCACTATAAGGCGAATTAAGAGAGAAAAGAAGAGTAAGAAGAAATATAAGACACCGGTCGCCACCATGGGAGAAAAGCCTGGCACCAGAGTGTTCAAGAAGTCTAGCCCCAACTGCAAGCTGACCGTGTACCTCGGCAAGCGGGACTTCGTGGACCACCTGGACAAAGTGGATCCCGTGGATGGCGTGGTGCTGGTGGATCCTGACTACCTGAAGGACCGGAAGGTGTTCGTGACCCTGACCTGCGCCTTCAGATACGGCAGAGAAGATCTGGACGTGCTGGGCCTGAGCTTCAGAAAGGACCTGTTTATCGCCACCTACCAGGCCTTTCCACCTGTGCCTAATCCTCCACGGCCTCCTACCAGACTGCAGGACAGACTGCTGAGAAAGCTGGGCCAGCACGCTCACCCCTTTTTCTTCACAATCCCTCAAAACCTGCCTTGCAGCGTGACACTGCAGCCCGGACCTGAGGATACAGGCAAAGCTTGCGGCGTGGACTTCGAGATCAGAGCCTTCTGTGCCAAGAGCCTGGAAGAGAAGTCCCACAAGAGAAACAGCGTGCGGCTGGTCATCAGAAAGGTGCAGTTCGCCCCTGAGAAGCCTGGACCTCAGCCTTCTGCCGAGACAACCAGACACTTCCTGATGAGCGACCGCAGCCTGCATCTGGAAGCCAGCCTCGACAAAGAGCTGTACTACCACGGCGAGCCCCTGAACGTCAACGTCCACGTGACCAACAACAGCACCAAGACCGTGAAAAAGATCAAGGTGTCCGTGCGGCAGTACGCCGACATCTGCCTGTTTAGCACAGCCCAGTACAAGTGCCCCGTGGCTCAGCTGGAACAGGACGATCAAGTGTCCCCTAGCAGCACCTTCTGCAAGGTGTACACAATCACCCCTCTGCTGAGCGACAACAGAGAGAAGAGAGGACTGGCCCTGGACGGCAAGCTGAAGCACGAGGATACAAACCTGGCCAGCAGCACCATCGTGAAAGAGGGCGCCAACAAAGAGGTGCTGGGCATCCTGGTGTCCTACAGAGTGAAAGTGAAGCTGGTGGTGTCCAGAGGCGGCGACGTTTCAGTGGAACTGCCCTTCGTGCTGATGCACCCCAAGCCTCACGATCACATCCCTCTGCCTAGACCTCAGAGCGCCGCTCCTGAAACAGATGTGCCCGTGGACACCAACCTGATCGAGTTCGACACCAACTACGCCACCGACGACGACATCGTGTTCGAGGACTTCGCCCGGCTGAGACTGAAGGGCATGAAGGACGACGACTACGACGATCAGCTTTGTGGCGGCGGAGGATCTGGCGGAGGTGGAAGCGGAGGCGGTGGATCTGAGAACCTGTACTTCCAGTTCCACAAGAGCGGCGCCTGGAAGCTGCCTGTGTCTCTGGTTAAGGGCGGCGGCGGCTCCGGCGGCGGCGGCTCCTACGTGCGGTTCGAGGTGCCCGAGGACATGCAGAACGAGGCCCTGAGCCTGCTGGAGAAGGTGCGGGAGAGCGGCAAGGTGAAGAAGGGCACCAACGAGACCACCAAGGCCGTGGAGCGGGGCCTTGCCAAGCTGGTGTACATCGCCGAGGACGTGGACCCCCCCGAGATCGTGGCCCACCTTCCTCTGCTGTGCGAGGAGAAGAACGTGCCCTACATCTACGTGAAGGAGAACCTGTACTTCCAGAGCAAGAACGACCTGGGCCGGGCCGTTGGCATCGAGGTGCCCTGCGCCAGTGCTGCCATCATCAACGAGGGCGAGCTGCGGAAGGAGCTGGGCAGCCTGGTGGAGAAGATCAAGGGCCTGCAGAAGTGAATCTAGACCTTCTGCGGGGCTTGCCTTCTGGCCATGCCCTTCTTCTCTCCCTTGCACCTGTACCTCTTGGTCTTTGAATAAAGCCTGAGTAGGAAAAAAAAAAAAAAAAAAAAAAAAAAAAAAAAAAAAAAAAAAAAAAAAAAAAAAAAAAAAAAAAAAAAAAAAAAAAAAAAAAAAAAAAAAAAAAAAAAAAAAAAAAAAAAAAAAAAAAAA]

- **2xScMS2(C)-IL1Ra-HiBiT**

T7 promoter (for CleanCap AG Reagent): 11-30

Stabilized 2xMS2-binding motif (C-variant): 51-117

Kozak sequence (including start codon): 125-133

Human IL-1Ra isoform II: 134-670

2xGSSG linker: 671-694

HiBiT: 695-727

Stop codon: 728-730

[CAGTGAATTGTAATACGACTCACTATAAGGTCAGATCCGCTAGCGGATCCGGGAGCAGGTGAGGATCACCCATCTGCCACGAGCGAGGTGAGGATCACCCATCTCGCTCGTGTTCCCACCGGTCGCCACCATGGCTCTGGCCGACCTGTATGAAGAAGGTGGCGGCGGAGGCGGAGAGGGCGAAGATAATGCCGACAGCAAAGAGACAATCTGCAGACCCAGCGGCCGGAAGTCCTCTAAGATGCAGGCCTTCAGAATCTGGGACGTGAACCAGAAAACCTTCTACCTGCGGAACAATCAGCTGGTGGCCGGCTATCTGCAGGGCCCCAATGTGAACCTGGAAGAGAAGATCGACGTGGTGCCCATCGAGCCCCACGCTCTGTTTCTTGGAATCCACGGCGGCAAGATGTGCCTGAGCTGTGTGAAGTCTGGCGACGAGACACGGCTGCAGCTGGAAGCCGTGAACATCACCGACCTGAGCGAGAACCGGAAGCAGGACAAGAGATTCGCCTTCATCAGAAGCGACAGCGGCCCCACCACAAGCTTTGAGTCTGCTGCTTGCCCTGGCTGGTTCCTGTGTACAGCCATGGAAGCCGACCAGCCTGTGTCTCTGACCAACATGCCTGACGAGGGCGTGATGGTCACCAAGTTCTACTTCCAAGAGGACGAGGGCAGCAGCGGCGGCAGCAGCGGCGTGAGCGGCTGGCGGCTGTTCAAGAAGATTAGCTGAATCTAGACCTTCTGCGGGGCTTGCCTTCTGGCCATGCCCTTCTTCTCTCCCTTGCACCTGTACCTCTTGGTCTTTGAATAAAGCCTGAGTAGGAAAAAAAAAAAAAAAAAAAAAAAAAAAAAAAAAAAAAAAAAAAAAAAAAAAAAAAAAAAAAAAAAAAAAAAAAAAAAAAAAAAAAAAAAAAAAAAAAAAAAAAAAAAAAAAAAAAAAAAA]

- **IL1Ra-HiBiT**

T7 promoter (for CleanCap AG Reagent): 11-30

Kozak sequence (including start codon): 79-87

Human IL-1Ra isoform II: 88-624

2xGSSG linker: 625-648

HiBiT: 649-681

Stop codon: 682-684

[CAGTGAATTGTAATACGACTCACTATAAGGCGAATTAAGAGAGAAAAGAAGAGTAAGAAGAAATATAAGACACCGGTCGCCACCATGGCTCTGGCCGACCTGTATGAAGAAGGTGGCGGCGGAGGCGGAGAGGGCGAAGATAATGCCGACAGCAAAGAGACAATCTGCAGACCCAGCGGCCGGAAGTCCTCTAAGATGCAGGCCTTCAGAATCTGGGACGTGAACCAGAAAACCTTCTACCTGCGGAACAATCAGCTGGTGGCCGGCTATCTGCAGGGCCCCAATGTGAACCTGGAAGAGAAGATCGACGTGGTGCCCATCGAGCCCCACGCTCTGTTTCTTGGAATCCACGGCGGCAAGATGTGCCTGAGCTGTGTGAAGTCTGGCGACGAGACACGGCTGCAGCTGGAAGCCGTGAACATCACCGACCTGAGCGAGAACCGGAAGCAGGACAAGAGATTCGCCTTCATCAGAAGCGACAGCGGCCCCACCACAAGCTTTGAGTCTGCTGCTTGCCCTGGCTGGTTCCTGTGTACAGCCATGGAAGCCGACCAGCCTGTGTCTCTGACCAACATGCCTGACGAGGGCGTGATGGTCACCAAGTTCTACTTCCAAGAGGACGAGGGCAGCAGCGGCGGCAGCAGCGGCGTGAGCGGCTGGCGGCTGTTCAAGAAGATTAGCTGAATCTAGACCTTCTGCGGGGCTTGCCTTCTGGCCATGCCCTTCTTCTCTCCCTTGCACCTGTACCTCTTGGTCTTTGAATAAAGCCTGAGTAGGAAAAAAAAAAAAAAAAAAAAAAAAAAAAAAAAAAAAAAAAAAAAAAAAAAAAAAAAAAAAAAAAAAAAAAAAAAAAAAAAAAAAAAAAAAAAAAAAAAAAAAAAAAAAAAAAAAAAAAAA]
